# Conserved molecular compatibility for MIZ1 function in cytokinin signaling in land plants

**DOI:** 10.64898/2026.10.12.758437

**Authors:** Adrijana Smoljan, Shubhajit Das, Dmitrii Vladimirstev, Anastasia Teplova, Marharyta Tkachenka, Wentao Yang, Mengting Lu, Eva Benková, Jiří Friml

## Abstract

Plant roots adjust their growth direction toward water through hydrotropism, a process that in *Arabidopsis thaliana* requires the endoplasmic reticulum (ER)-associated protein MIZU-KUSSEI1 (MIZ1). Nonetheless, MIZ1 orthologues occur throughout land plants, including lineages lacking true roots, raising questions about the ancestral function of this protein family. Here, we investigated the evolutionary conservation and functional diversification of MIZ1-like proteins across land plants. MIZ1 orthologues from evolutionarily distant land plants failed to restore hydrotropism or cytokinin sensitivity in the Arabidopsis *miz1* mutant, despite conserved protein structure and ER localization. In the liverwort *Marchantia polymorpha*, loss of *MpMIZ1a* led to altered cytokinin sensitivity and cytokinin-mediated development in both the sporophyte and gametophyte. These phenotypes and gene expression analyses imply a conserved role of MIZ1 in cytokinin-related processes. Despite this functional similarity, protein– protein docking and bimolecular fluorescence complementation revealed species-specific interactions between MIZ1 proteins and their cognate cytokinin histidine kinase receptors in Arabidopsis and Marchantia. Together, our findings indicate that MIZ1 proteins represent ancient land plant regulators of cytokinin-dependent development whose molecular compatibility and developmental outputs have diverged during evolution. We propose that the role of MIZ1 in root hydrotropism represents a derived recruitment of this ancestral module into root-specific environmental response pathways.

## Introduction

Water availability is a major determinant of plant growth and development and varies widely across soils and environments (Kramer and Boyer, 1995; Passioura, 2007). Plants respond to such variability through directional growth responses, including tropisms that guide organ growth in response to environmental gradients (Gilroy and Masson, 2008; Miyazawa and Takahashi, 2020). Among these, root hydrotropism, the directional growth of roots toward higher water availability, contributes to drought adaptation (Kiss, 2007; Robbins and Dinneny, 2015; Miyazawa and Takahashi, 2020).

In addition to primary root bending, plants adjust root system architecture in response to local water availability through processes such as hydropatterning and suppression of lateral root formation upon loss of water contact (Robbins and Dinneny, 2015; Giehl and von Wirén, 2018). Together, these responses highlight the importance of water-related cues in shaping root development (Orman-Ligeza et al., 2018). Despite growing insight into these responses, the mechanisms by which plants perceive water-related signals and convert them into directional growth responses are still being actively defined (Miyazawa and Takahashi, 2020).

Phytohormones play central roles in coordinating plant growth and environmental responses, including tropisms. Among these, auxin has long been established as a key regulator of several tropisms, including gravitropism and phototropism, through asymmetric distribution and differential growth responses (Han et al., 2021; Li et al., 2022). However, root hydrotropism differs from classical auxin-mediated tropisms and instead relies strongly on abscisic acid signaling, calcium dynamics, and cortex-based growth regulation (Takahashi et al., 2009; Dietrich et al., 2017; Shkolnik et al., 2018).

Genetic analyses in *Arabidopsis thaliana* have established hydrotropism as a tropic response that can be genetically uncoupled from gravitropism (Kobayashi et al., 2007). Forward genetic screens identified mutants defective in hydrotropism but largely normal in gravitropic responses, including *mizu-kussei1* (*miz1*) and *miz2* (Kobayashi et al., 2007; Miyazawa et al., 2009). *MIZ1* encodes a protein characterized by the MIZ1 domain, formerly DUF617 (Kobayashi et al., 2007). MIZ1 contributes to hydrotropic signaling by enabling a slow, long-distance Ca²⁺ signal required for root water tracking, in part through regulation of endoplasmic reticulum Ca²⁺ transport (Shkolnik et al., 2018). Consistent with this role, *MIZ1* expression is induced by osmotic stress and abscisic acid (Moriwaki et al., 2012), and MIZ1 mediates drought-triggered inhibition of root gravitropism during hydrotropism (Zhang et al., 2025).

Plant hormone cytokinin has also emerged as an important regulator of hydrotropic responses (Chang et al., 2019). Cytokinins regulate a broad range of developmental processes, such as cell division, organ growth, and senescence (Werner and Schmülling, 2009). Cytokinin signaling is mediated by a multistep two-component system involving CHASE domain-containing histidine kinase receptors, histidine-containing phosphotransfer proteins, and response regulators (Hwang et al., 2012; Kieber and Schaller, 2014). Type-B response regulators act as transcription factors that activate cytokinin-responsive genes, whereas type-A response regulators function as negative regulators that attenuate cytokinin signaling (Hwang et al., 2012; Kieber and Schaller, 2014). Genetic analyses in Arabidopsis have demonstrated essential roles for both classes of response regulators in multiple developmental contexts, including water-related growth responses (Werner and Schmülling, 2009).

Phylogenetic analyses indicate that the MIZ1 domain is specific to land plants and that many species encode multiple *MIZ1*-like genes, suggesting an early emergence followed by lineage-specific diversification (Kobayashi et al., 2007; Fujii et al., 2018). In Arabidopsis, MIZ1 proteins play a critical role in hydrotropic root bending. Notably, land plant species that lack true roots as water-absorbing organs also encode MIZ1 domain-containing proteins, suggesting that the ancestral function of MIZ1 proteins may not have been restricted to root hydrotropism (Miyazawa and Takahashi, 2020). However, experimental evidence addressing the functional conservation or divergence of *MIZ1*-like genes across major plant lineages is lacking.

*Marchantia polymorpha* is an early diverging land plant model with minimal genetic complexity that enables the investigation of ancestral protein functions and basic signaling mechanisms in land plant evolution (Ishizaki et al., 2016; Bowman et al., 2017). In contrast to Arabidopsis, which encodes 10 type-A and 11 type-B response regulators for cytokinin signaling, *M. polymorpha* contains a single gene for each type-A and type-B response regulator, providing a simplified genetic framework for studying cytokinin signaling (Aki et al., 2019). Cytokinin signaling has been implicated in key developmental processes in Marchantia, including thallus patterning and organ formation, yet how cytokinin-regulated pathways intersect with water-related growth responses in bryophytes remains unclear (Aki et al., 2019).

In this study, we investigate the evolutionary conservation and diversification of *MIZ1*-like genes across land plants using cross-species functional analyses and genetic approaches in a bryophyte model system.

## Results

### MIZ1-like proteins are conserved throughout land plants and localize to the endoplasmic reticulum

To investigate the evolutionary distribution of MIZ1-like proteins, we identified homologs using the Arabidopsis MIZ1 protein sequence as a query and performed a phylogenetic analysis using representative sequences from major land plant lineages. *MIZ1*-like genes were identified in bryophytes, lycophytes, gymnosperms, and angiosperms but were absent from green algae (Figure 1A). The recovered sequences formed 15 distinct clades with moderate to strong bootstrap support, indicating that MIZ1-like proteins originated early during land plant evolution and subsequently diversified. Structural predictions generated using AlphaFold2 revealed a conserved overall protein fold among representative MIZ1 homologs despite substantial sequence divergence (Figure S1). Structural superposition using the PyMOL plugin CEalign demonstrated overall structural similarity to *A. thaliana* MIZ1, with RMSD values of 1.83 Å for *M. polymorpha* MIZ1 (MpMIZ1), 2.66 Å for *Amborella trichopoda* MIZ1 (AmTrMIZ1), and 2.86 Å for *Physcomitrium patens* MIZ1 (PpMIZ1) (Supplementary Table 1).

**Figure 1.**
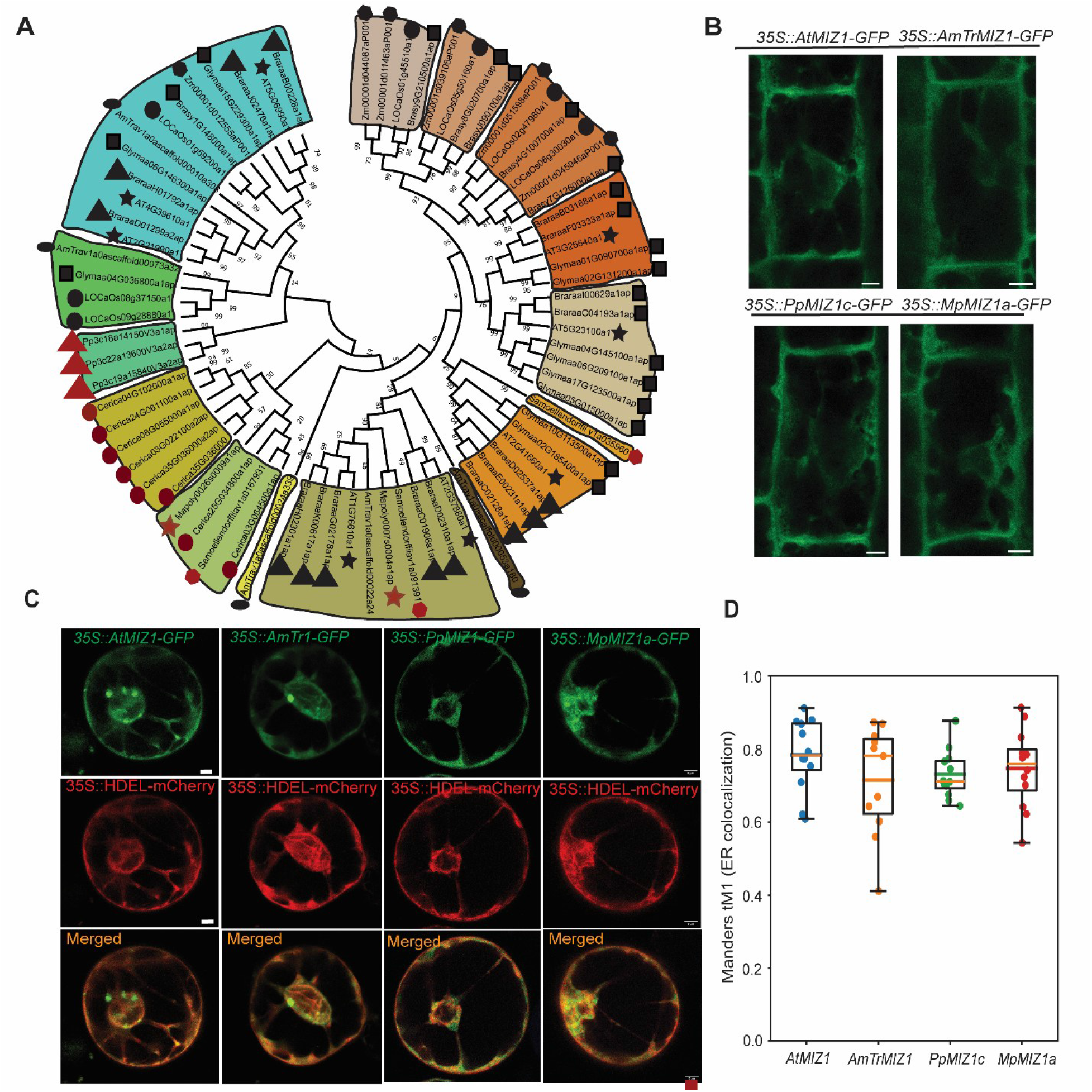
Evolutionarily conserved endoplasmic reticulum localization of MIZ1 proteins across land plants. (A) Maximum-likelihood phylogenetic analysis of MIZ1 proteins across land plants recovered 15 distinct clades with moderate to strong bootstrap support. Red symbols indicate MIZ1 homologs from non-seed land plants, including bryophytes and lycophytes, whereas black symbols indicate homologs from seed plants. Numbers at nodes represent bootstrap support values, and the scale bar indicates amino acid substitutions per site. (B) Representative confocal images of GFP-tagged MIZ1 proteins expressed in Arabidopsis roots, including *35S::AtMIZ1-GFP* in the *miz1* mutant background, *35S::AmTrMIZ1f-GF*P from *Amborella trichopoda*, *35S::PpMIZ1c-GFP* from *Physcomitrium patens*, and *35S::MpMIZ1a-GFP* from *Marchantia polymorpha*. (C) Co-localization of GFP-tagged MIZ1 proteins with the endoplasmic reticulum marker HDEL-mCherry in *Nicotiana benthamiana* protoplasts. GFP fluorescence is shown in green, mCherry fluorescence in magenta, and merged images are shown on the right. (D) Quantification of co-localization using Manders’ overlap coefficient (tM1) revealed strong co-localization of all examined MIZ1 homologs with the ER marker and no significant differences among proteins (Kruskal–Wallis test, H = 1.28, p = 0.734; Dunn’s post hoc test with Bonferroni correction, p > 0.05 for all pairwise comparisons). Points represent individual cells; bars indicate mean ± SD.

Arabidopsis MIZ1 is an endoplasmic reticulum (ER)-associated protein localized to the cytoplasmic face of the ER membrane (Yamazaki et al., 2012). To determine whether this subcellular localization is conserved among MIZ1 homologs from diverse plant lineages, GFP-tagged versions of *A. thaliana* MIZ1, *A. trichopoda* MIZ1f, *P. patens* MIZ1c, and *M. polymorpha* MIZ1a were expressed in the Arabidopsis *miz1* background and examined by confocal microscopy. All fusion proteins displayed a characteristic reticulate intracellular fluorescence pattern consistent with ER localization (Figure 1B).

To further validate ER association, the GFP-tagged proteins were transiently co-expressed with the ER marker HDEL-mCherry in Arabidopsis protoplasts. For all MIZ1 homologs tested, GFP fluorescence extensively overlapped with the HDEL-mCherry signal (Figure 1C). Quantitative colocalization analysis using Manders’ overlap coefficient (tM1) revealed high levels of overlap between each MIZ1 homolog and the ER marker, with no significant differences among the examined proteins (Figure 1D). These observations indicate that MIZ1 homologs from diverse land plant lineages share conserved localization to the endoplasmic reticulum.

### *MIZ1* homologs from early diverging land species fail to complement development of Arabidopsis *miz1*

To investigate whether MIZ1 homologs from evolutionarily distant land plant lineages can functionally substitute for Arabidopsis MIZ1, we expressed *A. trichopoda* MIZ1f (*35S::AmTrMIZ1f-GFP*), *P. patens* MIZ1c (*35S::PpMIZ1c-GFP*), and *M. polymorpha* MIZ1a (*35S::MpMIZ1a-GFP*) under the control of the constitutive 35S promoter in the Arabidopsis *miz1* background. Arabidopsis MIZ1 (*35S::AtMIZ1-GFP*) was used as a positive control. These are the same GFP-tagged transgenic lines used for the localization analyses shown in Figure 1. Additional homologs from *Selaginella moellendorffii* (*35S::SmMIZ1-GFP*), *P. patens* (*35S::PpMIZ1d-GFP*), and *M. polymorpha* (*35S::MpMIZ1b-GFP*) were analyzed in parallel and are presented in Supplementary Figure S1.

Hydrotropic responses were analyzed using a split medium consisting of control agar on one side and water-stress medium containing glycerol on the other side to generate a water potential gradient. While wild-type seedlings exhibited strong root bending in response to the gradient, *miz1* seedlings displayed a severely reduced hydrotropic response (Figure 2A,B). Expression of *AtMIZ1* restored hydrotropic bending in the *miz1* background. In contrast, expression of *MIZ1* homologs from *A. trichopoda*, *P. patens*, and *M. polymorpha* failed to rescue the hydrotropic defect, and the corresponding transgenic lines exhibited bending angles comparable to those of *miz1* (Figure 2A,B). A similar lack of complementation was observed for *SmMIZ1*, *PpMIZ1d*, and *MpMIZ1b*, which also failed to restore hydrotropic bending in the *miz1* background (Supplementary Figure S2).

**Figure 2.**
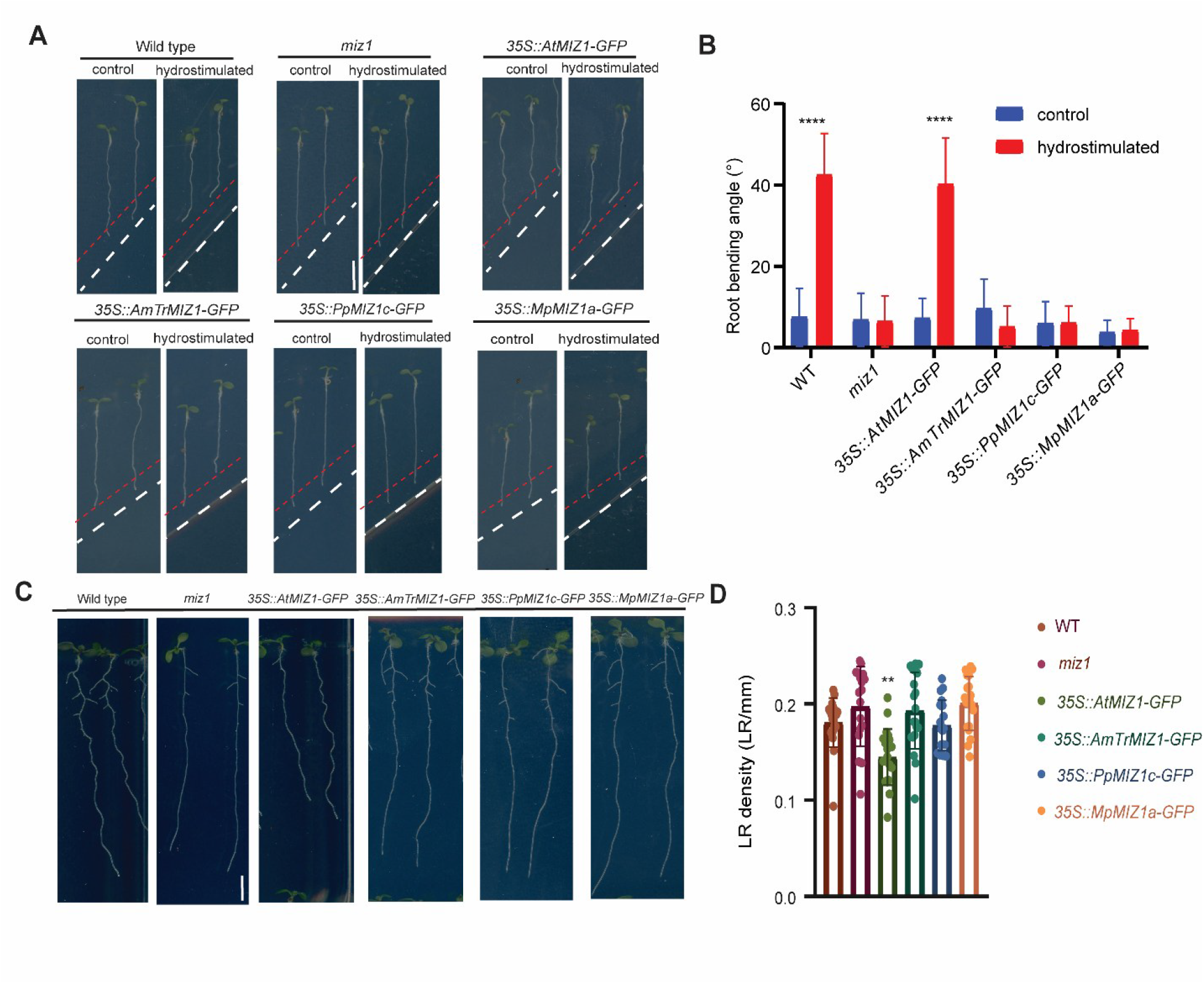
Functional analysis of MIZ1 homologs in the Arabidopsis *miz1* mutant background. (A) Representative images of the hydrotropic response of WT, *miz1*, *35S::AtMIZ1-GFP*, *35S::AmTrMIZ1f-GFP*, *35S::PpMIZ1c-GFP*, and *35S::MpMIZ1a-GFP* seedlings grown on a split medium consisting of agar and water-stress medium. Scale bar = 0.5 cm. (B) Quantification of root bending angles under control and hydrostimulated conditions. Data were analyzed by two-way ANOVA followed by Sidak’s multiple comparisons test. (C) Representative images of WT, *miz1*, *35S::AtMIZ1-GFP*, *35S::AmTrMIZ1f-GFP*, *35S::PpMIZ1c-GFP*, and *35S::MpMIZ1a-GFP* seedlings used for lateral root density analysis. Scale bar = 0.5 cm. (D) Quantification of lateral root density. Data were analyzed by one-way ANOVA followed by Dunnett’s multiple comparisons test. Bars represent mean ± SD; n = 20 seedlings per genotype.

We next quantified lateral root density in the same set of transgenic lines (Figure 2C,D). Previous studies demonstrated that although the *miz1* mutant does not exhibit a pronounced defect in lateral root development under normal growth conditions, overexpression of Arabidopsis *MIZ1* suppresses lateral root formation and results in reduced lateral root density (Moriwaki et al., 2012). Consistent with these findings, seedlings expressing *35S::AtMIZ1-GFP* exhibited a significant reduction in lateral root density compared with wild-type plants. In contrast, lines expressing *MIZ1* homologs from *A. trichopoda*, *P. patens*, and *M. polymorpha* displayed lateral root densities comparable to those of *miz1* and did not reproduce the characteristic overexpression phenotype of Arabidopsis *MIZ1* (Figure 2D). Likewise, *SmMIZ1*, *PpMIZ1d*, and *MpMIZ1b* lines were indistinguishable from *miz1* with respect to lateral root density (Supplementary Figure S2).

To determine whether the lack of complementation observed in agar-based assays also occurred under more natural growth conditions, we analyzed root growth in soil using a rhizotron system that generates stable moisture gradients (Supplementary Figure S3). Wild-type plants developed asymmetric root system architectures biased toward regions of higher soil moisture, whereas *miz1* plants failed to exhibit this root development asymmetry. Expression of *AtMIZ1* restored this response in the *miz1* background, while lines expressing *MIZ1* homologs from *A. trichopoda*, *P. patens*, and *M. polymorpha* remained indistinguishable from *miz1* (Supplementary Figure S3).

Together, these results indicate that MIZ1 homologs from bryophytes, lycophytes, and a basal angiosperm are unable to complement the hydrotropic and hydropatterning defects of the Arabidopsis *miz1* mutant and do not reproduce the lateral root phenotype associated with Arabidopsis *MIZ1* overexpression.

### *MIZ1* homologs from early diverging land plants fail to restore cytokinin sensitivity in Arabidopsis *miz1*

Recent work demonstrated that MIZ1 modulates cytokinin signaling during root development through interactions with cytokinin receptors and that *miz1* mutants exhibit reduced sensitivity to cytokinin, whereas *MIZ1* overexpression enhances cytokinin-mediated inhibition of root growth (Fu et al., 2025). To further investigate whether *MIZ1* orthologues from evolutionarily distant land plants can substitute for Arabidopsis MIZ1, we examined cytokinin sensitivity using exogenous zeatin treatment.

The same *35S::MIZ1-GFP* transgenic lines used for the localization and complementation analyses were used for cytokinin response assays. Wild-type, *miz1*, *35S::AtMIZ1-GFP*, and transgenic lines expressing *MIZ1* homologs from *A. trichopoda*, *P. patens*, and *M. polymorpha* were grown on medium supplemented with 1 μM zeatin. Representative seedling phenotypes are shown in Figure 3A, and quantification of primary root length is presented in Figure 3B. As expected, *miz1* seedlings displayed reduced sensitivity to cytokinin treatment, whereas expression of *AtMIZ1* restored cytokinin responsiveness, resulting in significantly shorter primary roots in the presence of zeatin. In contrast, transgenic lines expressing *MIZ1* homologs from *A. trichopoda*, *P. patens*, and *M. polymorpha* exhibited primary root lengths comparable to those of *miz1* and did not show restored cytokinin sensitivity (Figure 3B).

**Figure 3.**
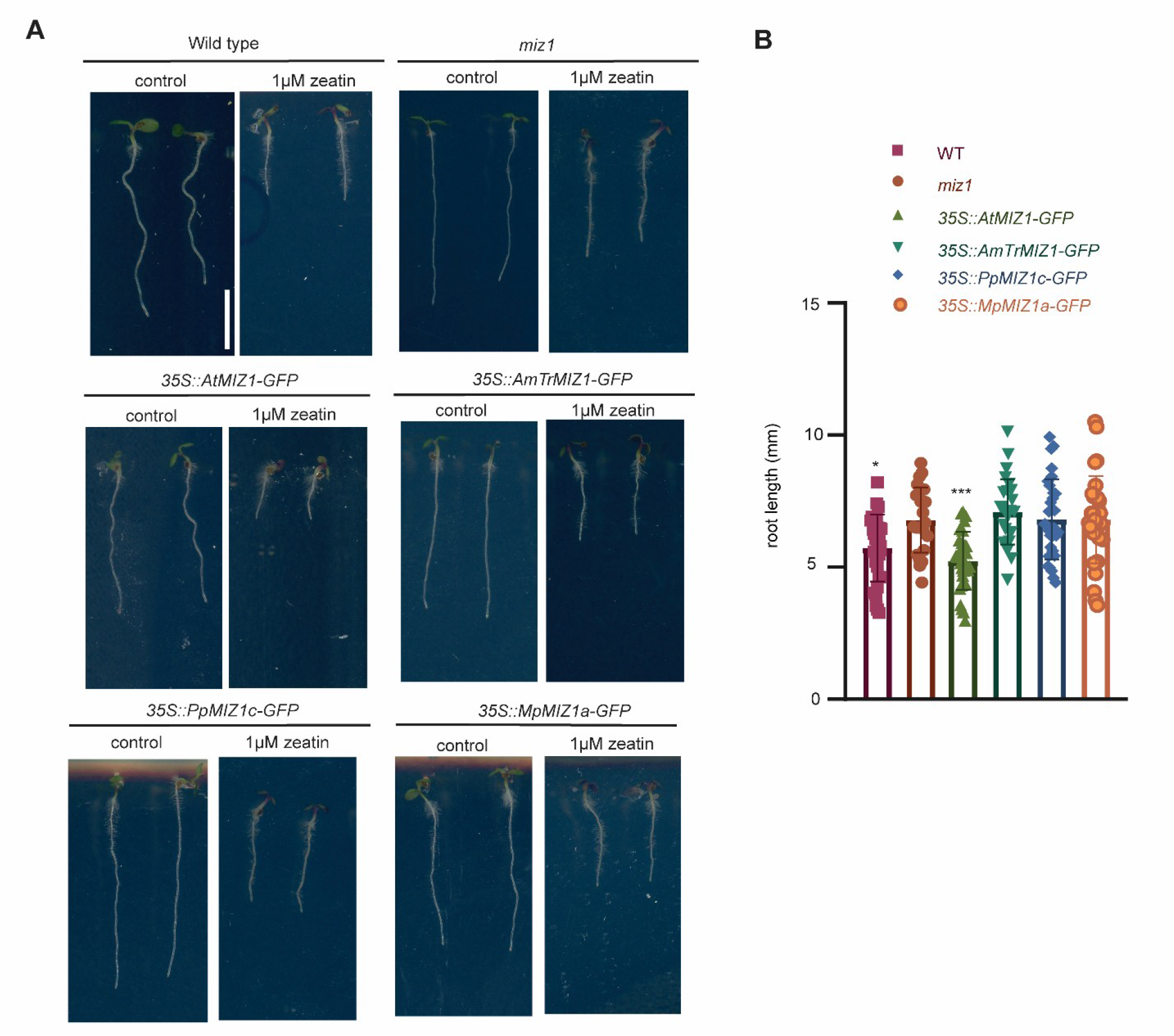
Response of MIZ1 transgenic lines to cytokinin treatment. (A) Representative images of WT, *miz1*, *35S::AtMIZ1-GFP*, *35S::AmTrMIZ1f-GFP*, *35S::PpMIZ1c-GFP*, and *35S::MpMIZ1a-GFP* seedlings grown on mock medium (left) or medium supplemented with 1 μM zeatin (right). Scale bar = 0.5 cm. (B) Quantification of primary root length of seedlings grown on medium supplemented with 1 μM zeatin. Data were analyzed by one-way ANOVA followed by Dunnett’s multiple comparisons test using *MIZ1* as the reference genotype. Bars represent mean ± SD and individual points represent biological replicates.

To determine whether this lack of complementation was a general feature of non-Arabidopsis *MIZ1* homologs, we also examined transgenic lines expressing homologs from *S. moellendorffii (SmMIZ1)*, *P. patens* (*PpMIZ1d*), and *M. polymorpha* (MpMIZ1b). All three lines displayed cytokinin responses indistinguishable from those of *miz1* and failed to restore sensitivity to zeatin treatment, similar to the homologs shown in Figure 3 (Supplementary Figure S4).

Together, these results show that MIZ1 homologs from evolutionarily distant land plants are unable to substitute for Arabidopsis MIZ1 in cytokinin-mediated root growth regulation. Notably, this lack of complementation was observed not only for homologs from bryophytes and lycophytes but also for the *A. trichopoda* homolog, despite its origin from a basal angiosperm. These findings suggest that the specific function of Arabidopsis MIZ1 in cytokinin-mediated root growth is not functionally interchangeable with MIZ1 homologs from the tested land plant lineages.

### MpMIZ1a regulates thallus growth and cytokinin homeostasis in Marchantia

Since early MIZ1 homologs failed to restore *Atmiz1* phenotypes, we wondered what could be the native functions of these homologs in early diverging land plant species. To explore the ancestral function of MIZ1 proteins, we generated loss-of-function CRISPR mutants and overexpression lines for the two *M. polymorpha MIZ1* genes, *MpMIZ1a* and *MpMIZ1b* in Tak-1 (wild-type male) background. Under mock conditions, both independent *Mpmiz1a* mutant alleles (*miz1a#1* and *miz1a#2*) exhibited hyponastic thallus growth and significantly reduced thallus area compared with Tak-1 (Figure 4A,B). In contrast, *Mpmiz1b* mutants (*miz1b#12* and *miz1b#13*), as well as the overexpression lines *pUBE2::MIZ1A-mNG* and *pUBE2::MIZ1B-mNG*, were comparable to wild-type plants. Quantification of thallus area confirmed that the growth defect was specific to the loss of *MpMIZ1a* (Figure 4B). The reduced thallus size of *Mpmiz1a* mutants was also observed in soil-grown plants at three weeks of age (Supplementary Figure S5). Furthermore, *Mpmiz1a* mutants produced an increased number of rhizoids relative to wild-type plants, whereas *Mpmiz1b* mutants did not display obvious alterations in rhizoid development (Supplementary Figure S6).

**Figure 4.**
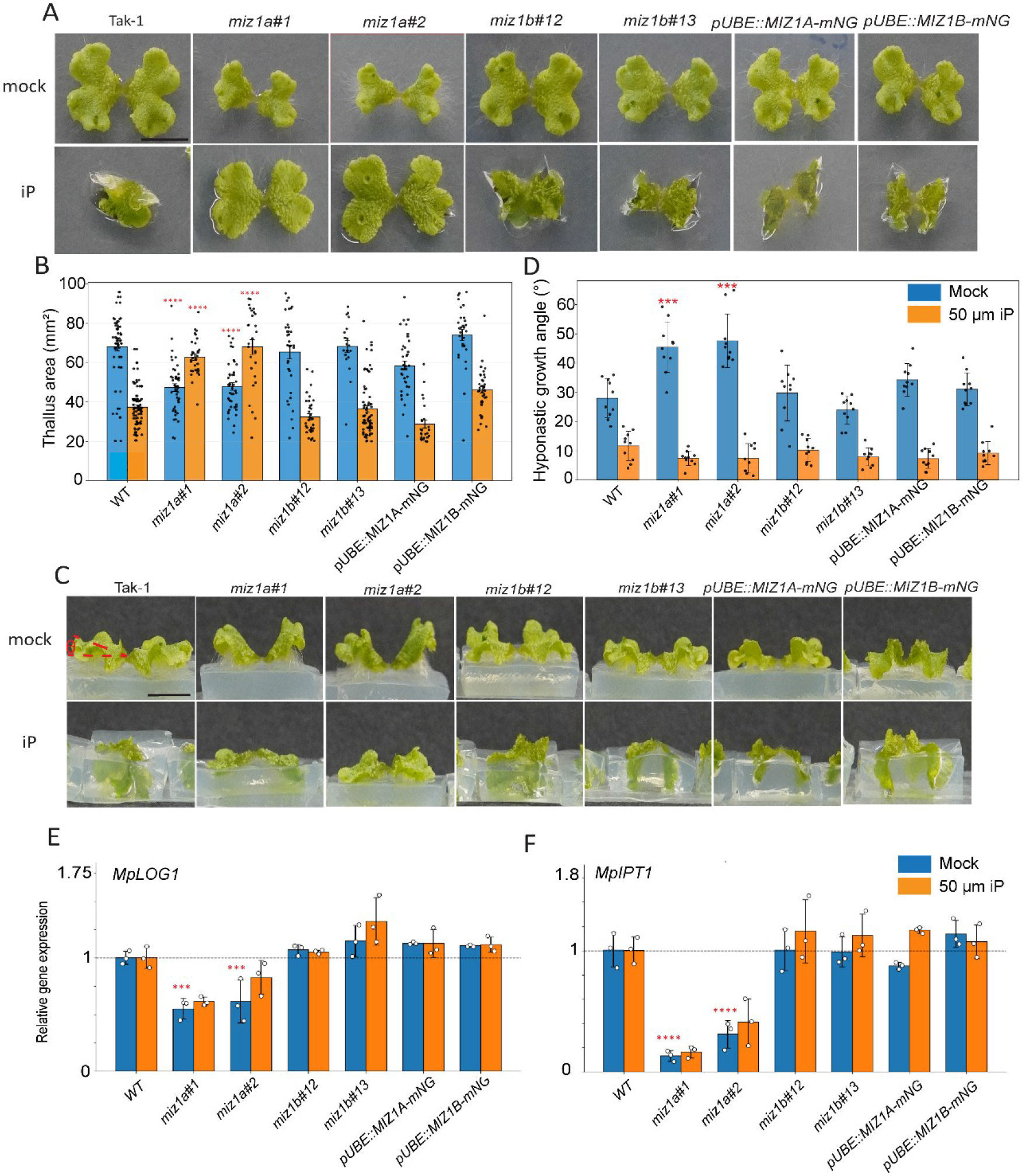
Cytokinin responses in *Marchantia polymorpha MIZ1A* and *MIZ1B* mutant and overexpression lines. (A) Representative images of Tak-1, *MIZ1a#1*, *MIZ1a#2*, *MIZ1b#12*, *MIZ1b#13*, *pUBE2::MIZ1A-mNG*, and *pUBE2::MIZ1B-mNG* thalli grown on mock medium (upper panel) or medium supplemented with 50 μM *N*⁶-isopentenyladenine (iP; lower panel). Scale bar = 0.5 cm. (B) Quantification of thallus area (mm²) of plants grown under mock and 50 μM iP conditions. Comparisons between genotypes and Tak-1 within each treatment were performed using unpaired Welch’s *t*-tests or Mann–Whitney *U* tests where appropriate. (C) Representative images showing hyponastic growth responses of Tak-1, *MIZ1a#1*, *MIZ1a#2*, *MIZ1b#12*, *MIZ1b#13*, *pUBE2::MIZ1A-mNG*, and *pUBE2::MIZ1B-mNG* grown on mock medium (upper panel) or medium supplemented with 50 μM iP (lower panel). Scale bar = 0.5 cm. (D) Quantification of hyponastic growth angle (°) under mock and 50 μM iP conditions. Comparisons between genotypes and Tak-1 within each treatment were performed using unpaired Welch’s *t*-tests or Mann–Whitney *U* tests where appropriate. (E) Relative expression of *MpLOG1* in Tak-1, *MIZ1a*#1, *MIZ1a*#2, *MIZ1b*#12, *MIZ1b*#13, pUBE2::*MIZ1A*-mNG, and pUBE2::*MIZ1B*-mNG plants grown under mock or 50 μM iP conditions, determined by quantitative RT-qPCR. Transcript levels were normalized to the reference gene (EF1 alpha) and expressed relative to Tak-1 within the corresponding treatment. Bars represent the mean ± SD, and individual points represent technical replicates. (F) Relative expression of *MpIPT1* in Tak-1, *MIZ1a*#1, *MIZ1a*#2, *MIZ1b*#12, *MIZ1b*#13, pUBE2::*MIZ1A*-mNG, and pUBE2::*MIZ1B*-mNG plants grown under mock or 50 μM iP conditions, determined by quantitative RT-qPCR. Transcript levels were normalized to the reference gene and expressed relative to Tak-1 within the corresponding treatment. Bars represent the mean ± SD.

Cytokinin signaling plays a central role in thallus development and organ formation in *M. polymorpha* (Aki et al., 2019). To examine the involvement of MpMIZ1 proteins in cytokinin-regulated growth, gemmae were grown on medium supplemented with 50 μM *N*6-isopentenyladenine (iP). Under cytokinin treatment, wild-type plants exhibited epinastic thallus bending and a reduction in thallus area relative to mock-grown controls (Figure 4A,B). In contrast, cytokinin treatment rescued the hyponastic growth phenotype of both *Mpmiz1a* mutant alleles. Both alleles displayed flat thallus growth and increased thallus area under iP treatment compared with their corresponding mock controls. In contrast, *Mpmiz1b* mutants and the overexpression lines responded similarly to wild-type Tak-1 (Figure 4A,B).

We next assessed growth orientation by measuring thallus growth angles. Under mock conditions, *Mpmiz1a* mutants exhibited increased upward bending angles compared with wild-type plants and the other genotypes analyzed (Figure 4C,D). Treatment with 50 μM iP reduced bending angles in all genotypes. In wild-type plants, *Mpmiz1b* mutants, and the overexpression lines, cytokinin induced downward thallus bending, resulting in growth into the medium. In *Mpmiz1a* mutants, cytokinin treatment suppressed the hyponastic growth phenotype and restored flat thallus growth but did not induce the pronounced epinastic growth observed in the other genotypes (Figure 4C). Together, these findings identify *MpMIZ1a* as the major MIZ1 homolog regulating thallus growth and cytokinin-dependent growth responses in *M. polymorpha*.

To further investigate whether the altered cytokinin responses of *Mpmiz1a* mutants are associated with changes in cytokinin biosynthesis or signaling, we examined the expression of genes involved in cytokinin metabolism together with a cytokinin-responsive marker. Cytokinin biosynthesis in land plants is primarily regulated by isopentenyltransferases (IPTs), which catalyze a rate-limiting step in cytokinin biosynthesis, and LONELY GUY (LOG) phosphoribohydrolases, which convert inactive cytokinin nucleotides into biologically active free-base forms (Kurakawa et al., 2007). We therefore quantified the expression of *MpIPT1*, *MpLOG1*, and the cytokinin-responsive type-A response regulator *MpRRA* following treatment with 50 μM iP (Figure 4E,F; Supplementary Figure S7).

Under mock conditions, both independent *Mpmiz1a* mutant alleles displayed consistently reduced expression of *MpIPT1* and *MpLOG1* compared with Tak-1, whereas *Mpmiz1b* mutants and the *MpMIZ1* overexpression lines showed transcript levels comparable to those of wild-type plants. Treatment with 50 μM iP did not significantly alter *MpIPT1* or *MpLOG1* expression in Tak-1 compared with mock conditions. Similarly, expression of both genes remained reduced in the *Mpmiz1a* mutant alleles relative to Tak-1, whereas *Mpmiz1b* mutants and the *MpMIZ1* overexpression lines maintained transcript levels comparable to wild type under iP treatment. In contrast, exogenous cytokinin strongly induced *MpRRA* expression in Tak-1, and a comparable induction was observed in both *Mpmiz1a* and *Mpmiz1b* mutants, as well as in the *MpMIZ1* overexpression lines (Supplementary Figure S7), indicating that cytokinin-responsive signaling remains functional in all genotypes.

Together, the reduced expression of *MpIPT1* and *MpLOG1* in *Mpmiz1a* mutants, combined with the retained induction of the cytokinin-responsive marker *MpRRA*, suggests that loss of *MpMIZ1a* primarily affects cytokinin homeostasis rather than abolishing cytokinin signal transduction. These findings are consistent with the reported role of Arabidopsis MIZ1 in modulating cytokinin homeostasis through the regulation of genes involved in cytokinin biosynthesis and catabolism (Fu et al., 2025) and suggest that *MpMIZ1a* similarly contributes to the maintenance of cytokinin homeostasis.

### MpMIZ1a is highly expressed in spores and regulates spore development

To gain insight into the developmental context of *MpMIZ1a* expression in *M. polymorpha*, we examined publicly available transcriptomic datasets through MarpolBase. These data indicated that *MpMIZ1a* is expressed in several tissues, including thalli and gemma cups, but reaches its highest expression levels in spores. Based on this expression pattern, we investigated MpMIZ1a protein accumulation during spore germination.

To visualize MpMIZ1a protein accumulation *in vivo*, we generated a translational reporter line expressing *MpMIZ1a-mNeonGreen* under the control of the native *MpMIZ1a* promoter and regulatory sequences. Reporter lines generated in the male Tak-1 background were crossed with female Tak-2 plants to obtain F1 spores, approximately 50% of which were expected to carry the mNeonGreen-tagged transgene. MpMIZ1a-mNeonGreen fluorescence was detected in developing spores using confocal microscopy. As expected, approximately half of the spores exhibited detectable mNeonGreen fluorescence. A strong MpMIZ1a-mNeonGreen signal was observed immediately after spore release (0 h), indicating high protein accumulation at this stage (Figure 5A). The fluorescence signal was substantially reduced after 24 h and was no longer detectable after 48 h of spore germination, indicating a decline in MpMIZ1a protein accumulation during early developmental progression (Figure 5A).

**Figure 5.**
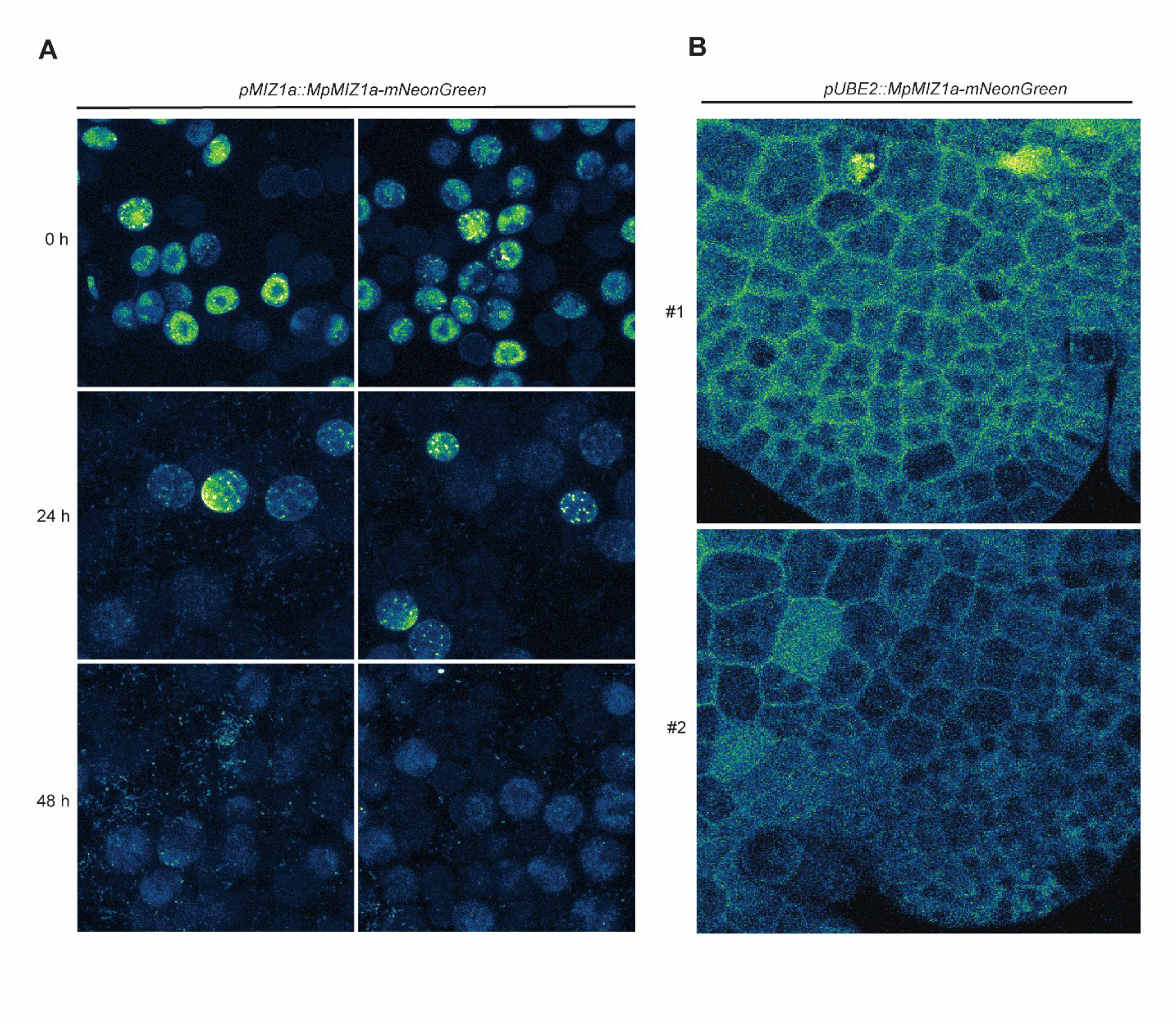
MpMIZ1a is highly expressed in spores. (A) Representative confocal images of *pMIZ1a::MIZ1a-mNG* spores at 0 h, 24 h, and 48 h after germination. MIZ1a-mNeonGreen fluorescence is detected in dormant spores and the signal progressively decreases during spore germination. (B) Representative confocal images of gemmae from two independent *pUBE2::MIZ1a-mNG* lines (#1 and #2). MIZ1a-mNeonGreen fluorescence is visible throughout the gemmae.

To determine whether MpMIZ1a protein also accumulates in vegetative tissues, where *Mpmiz1a* mutants exhibit altered cytokinin responses, we examined MpMIZ1a-mNeonGreen fluorescence in gemmae of the native-promoter reporter lines. Unlike in spores, where strong fluorescence was detected, no detectable signal was observed in gemmae. To investigate whether MpMIZ1a could accumulate in vegetative tissues under constitutive expression, we examined two independent *pUBE2::MIZ1A-mNeonGreen* overexpression lines. Despite expression driven by the ubiquitin promoter, only weak fluorescence was detected in gemmae (Figure 5B), in marked contrast to the strong accumulation observed in spores expressing the native-promoter reporter. Furthermore, no detectable fluorescence was observed in *pUBE2::MIZ1B-mNG* gemmae under the same imaging conditions (Supplementary Figure S8). Together, these observations indicate that MpMIZ1a protein accumulation differs markedly between reproductive and vegetative tissues, with high levels detected in spores but only weak or undetectable fluorescence in gemmae, even under constitutive expression.

As the translational reporter lines revealed high MpMIZ1a protein accumulation in early spores, we asked whether MpMIZ1 proteins contribute to spore development. To this end, we examined spore formation in *miz1a* and *miz1b* mutant lines. Both mutants were generated in the Tak-1 background and developed normal gametangiophores upon induction with far-red light. However, following crossing with wild-type Tak-2 females, *miz1b* mutants developed sporangia comparable to those of wild-type plants, whereas crosses with *miz1a* mutants showed a severe defect in spore formation, with sporangia either absent or their formation strongly reduced in the analyzed plants (Supplementary Figure S9). These observations indicate that *MpMIZ1a*, but not *MpMIZ1b*, is required for normal sporophyte development in *M. polymorpha*.

Together, these findings identify MpMIZ1a as a developmentally regulated MIZ1 protein with highest accumulation during the spore stage and reveal an essential role for *MpMIZ1a* in spore formation. The decline in MpMIZ1a protein abundance during germination further suggests that its function may be particularly important during early developmental transitions. This expression pattern is consistent with the conserved role of cytokinin in regulating reproductive development and early developmental transitions in land plants, including embryo development, seed formation, and germination in Arabidopsis (Riefler et al., 2006; Bartrina et al., 2011), supporting a role for MpMIZ1a in cytokinin-dependent developmental processes during the reproductive phase of the Marchantia life cycle.

### MIZ1 interaction with cytokinin receptors is conserved but species specific

In Arabidopsis, it was recently reported that MIZ1 physically interacts with the cytokinin receptors AHK2, AHK3, and AHK4 and that these interactions are required for MIZ1-mediated regulation of cytokinin signaling and hydrotropism (Fu et al., 2025). The inability of liverwort, lycophyte, and basal angiosperm MIZ1 homologs orthologues to restore cytokinin sensitivity in the Arabidopsis *miz1* mutant prompted us to investigate whether differences in cross-species MIZ1–AHK receptor interactions could underlie the lack of functional complementation.

Although AtMIZ1–AtAHK interactions were reported using yeast two-hybrid and co-immunoprecipitation assays (Fu et al., 2025), the molecular interfaces responsible for these interactions and the exact amino acid residues involved in binding remain unknown. To identify potential interaction sites, we performed protein– protein docking analyses between AtMIZ1 and AtAHK2 using HDOCK (Yan et al., 2017). Structural models of both proteins were obtained using AlphaFold predictions (Jumper et al., 2021). The predicted AtMIZ1–AtAHK2 complex revealed a discrete interaction interface involving multiple residues from both proteins (Figure 6A). Analysis of the docking model identified several potential hydrogen bonds and salt bridges that may contribute to complex formation (Figure 6B,C). Putative interacting residues were identified based on intermolecular distances of 2.5–3.0 Å.

**Figure 6.**
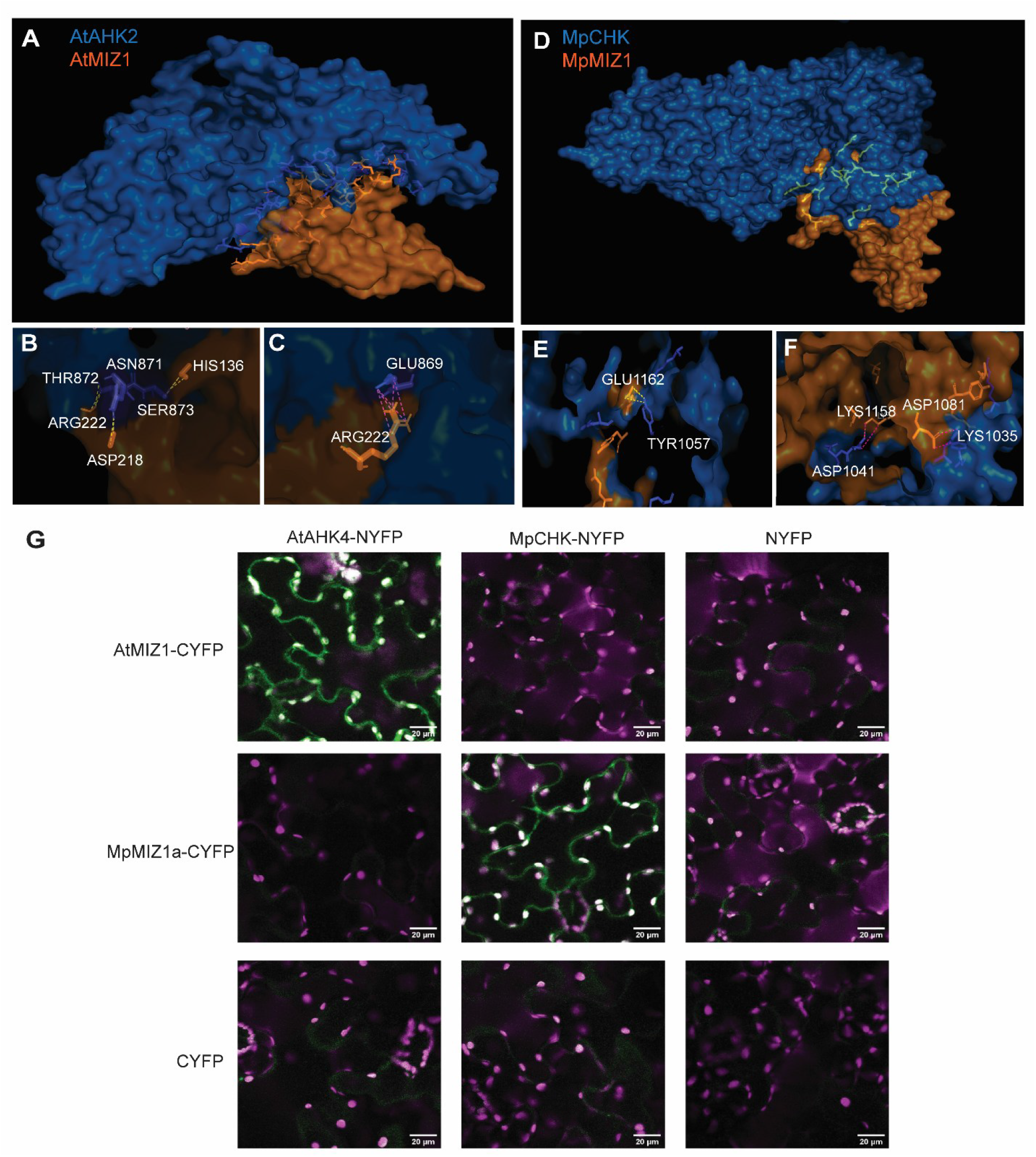
Predicted interactions between MIZ1 proteins and cytokinin receptors and validation by bimolecular fluorescence complementation. (A) Predicted protein-protein interaction model between *Arabidopsis thaliana* MIZ1 and the cytokinin receptor AtAHK2 generated using HDOCK. The AtMIZ1 structure was obtained using AlphaFold co-fold prediction. Putative interface residues are highlighted as light blue sticks. (B) Predicted hydrogen bonds between AtMIZ1 and AtAHK2. Residues involved in hydrogen bond formation are indicated, and hydrogen bonds are shown as yellow dashed lines. (C) Predicted salt bridges at the AtMIZ1-AtAHK2 interaction interface. Residues participating in salt bridge formation are indicated, and salt bridges are shown as pink dashed lines. (D) Predicted protein-protein interaction model between *Marchantia polymorpha* MpMIZ1a and the cytokinin receptor MpCHK generated using HDOCK. Putative interface residues are highlighted as light blue sticks. (E) Predicted hydrogen bonds between MpMIZ1a and MpCHK. Residues involved in hydrogen bond formation are indicated, and hydrogen bonds are shown as yellow dashed lines. (F) Predicted salt bridges at the MpMIZ1a-MpCHK interaction interface. Residues participating in salt bridge formation are indicated, and salt bridges are shown as pink dashed lines. Putative interacting residues shown in panels B, C, E, and F were identified based on intermolecular distances of 2.5–3.0 Å. (G) Bimolecular fluorescence complementation (BiFC) analysis of interactions between MIZ1 proteins and cytokinin histidine kinases in *Nicotiana benthamiana* leaves. Representative confocal images following transient co-expression of AtMIZ1-CYFP or MpMIZ1a-CYFP with AtAHK4-NYFP or MpCHK-NYFP are shown. Reconstitution of YFP fluorescence (green) indicates protein-protein interaction. Strong BiFC signals were detected for the cognate AtMIZ1-AtAHK4 and MpMIZ1a-MpCHK pairs, whereas little or no YFP fluorescence was observed for the reciprocal cross-species combinations. Co-expression with the corresponding NYFP- or CYFP-only constructs served as negative controls. Chlorophyll autofluorescence is shown in magenta. Scale bars, 20 µm.

We next asked whether a similar interaction could be predicted between the Marchantia proteins MpMIZ1a and MpCHK, the Marchantia cytokinin histidine kinase (Aki et al., 2019). Docking analyses generated a stable MpMIZ1a–MpCHK interaction model, with a predicted interface distinct from, but structurally analogous to, that observed for the Arabidopsis components (Figure 6D). Similar to the AtMIZ1–AtAHK2 model, several candidate hydrogen bonds and salt bridges were identified at the predicted MpMIZ1a– MpCHK interaction surface (Figure 6E–F), suggesting that MpMIZ1a may also have the capacity to associate with cytokinin receptors.

To experimentally assess interactions between MIZ1 proteins and cytokinin histidine kinases, we performed bimolecular fluorescence complementation (BiFC) assays in *Nicotiana benthamiana* leaves. AtMIZ1 or MpMIZ1a fused to the C-terminal half of YFP (CYFP) were co-expressed with either AtAHK4 or MpCHK fused to the N-terminal half of YFP (NYFP). Strong reconstitution of YFP fluorescence was observed for the cognate AtMIZ1–AtAHK4 and MpMIZ1a–MpCHK combinations (Figure 6G), supporting an interaction between MIZ1 proteins and cytokinin histidine kinases in both Arabidopsis and Marchantia. By contrast, little or no YFP fluorescence was detected for the reciprocal cross-species combinations, AtMIZ1– MpCHK and MpMIZ1a–AtAHK4. Co-expression with the corresponding NYFP- or CYFP-only constructs similarly showed no detectable BiFC signal.

Together, the docking predictions and BiFC analyses support a conserved association between MIZ1 proteins and cytokinin histidine kinases, while indicating that the compatibility of these interactions has diverged between Arabidopsis and Marchantia. In particular, MpMIZ1a interacted with its cognate cytokinin histidine kinase MpCHK but showed little or no detectable interaction with the Arabidopsis receptor AtAHK4. Conversely, AtMIZ1 interacted with AtAHK4 but not detectably with MpCHK. This species-specific interaction pattern provides a potential mechanistic explanation for the inability of MpMIZ1a to restore cytokinin sensitivity and hydrotropic responses in the Arabidopsis *miz1* mutant. Thus, although an association between MIZ1 proteins and cytokinin signaling components appears to be conserved across land plants, molecular compatibility between the respective interaction partners have diverged during evolution.

## Discussion

The evolutionary origin of hydrotropism remains poorly understood despite the identification of key regulators in Arabidopsis. MIZ1 is particularly intriguing because it is required for hydrotropic root bending, yet homologs are present throughout land plants, including lineages that lack true roots. This prompted a question of whether MIZ1 originally evolved as a hydrotropism regulator or whether this function emerged later through recruitment of an ancestral signaling component into root-specific developmental pathways. By combining cross-species complementation with subcellular localization, protein interaction analyses, and functional characterization in early diverging land plant *Marchantia polymorpha*, our results support the latter scenario and reveal substantial functional diversification of MIZ1 proteins during land plant evolution.

### MIZ1 originated early in land plant evolution but its hydrotropic function is not broadly conserved

Our phylogenetic analysis identified MIZ1-like proteins throughout land plants but not in green algae, consistent with previous analyses suggesting that the MIZ1 domain is a land plant innovation (Kobayashi et al., 2007; Fuji et al., 2018). The presence of MIZ1 homologs in bryophytes, lycophytes, gymnosperms, and angiosperms indicates that this protein family has been maintained throughout land plant diversification.

Despite this broad conservation, homologs from Marchantia, Physcomitrium, Selaginella, and Amborella failed to restore hydrotropism or cytokinin sensitivity in the Arabidopsis *miz1* mutant. The inability of even the basal angiosperm Amborella homolog to substitute for AtMIZ1 suggests that functional divergence is not restricted to the evolutionary transition between nonvascular and vascular plants or to the origin of roots. Instead, specific molecular features required for MIZ1-mediated hydrotropism in Arabidopsis appear to have evolved later within seed plant or angiosperm evolution. Thus, the evolutionary conservation of MIZ1 proteins does not correspond to conservation of their hydrotropic function.

### Conserved molecular properties despite functional divergence

The lack of functional complementation contrasts with the conservation of other molecular properties. All tested MIZ1 homologs localized to the endoplasmic reticulum, consistent with previous observations for Arabidopsis MIZ1 (Yamazaki et al., 2012), indicating that ER association represents an evolutionarily conserved feature of the MIZ1 protein family. However, this conserved localization was insufficient to confer AtMIZ1-like function, suggesting that functional divergence must involve other molecular properties.

Recent work identified Arabidopsis MIZ1 as a regulator of both cytokinin signaling and homeostasis. MIZ1 directly interacts with the cytokinin receptors AHK2, AHK3, and AHK4 and affects cytokinin-responsive gene expression, while also influencing the expression of genes involved in cytokinin biosynthesis and catabolism (Fu et al., 2025). Our interaction analyses suggest that the association between MIZ1 proteins and cytokinin receptors may itself represent a conserved feature as Marchantia MpMIZ1a interacted with the cytokinin receptor MpCHK. This suggests that association of MIZ1 proteins with cytokinin regulation evolved very early in MIZ1 evolution.

Nonetheless, the cross-species interactions between MpMIZ1a and AtAHK2 or AtMIZ1 and MpCHK were not detected. This species-specific compatibility provides a possible molecular explanation for the inability of MpMIZ1a to complement the Arabidopsis *miz1* mutant as Marchantia MpMIZ1a is unable to engage correctly with the Arabidopsis cytokinin signaling machinery. The conservation of receptor association within species, together with the loss of detectable interaction across species, therefore suggests co-divergence of MIZ1 proteins and components of their cytokinin-related molecular environment. Further biochemical analyses will be required to determine the molecular determinants underlying this species specificity.

### MpMIZ1a links MIZ1 proteins to cytokinin-dependent development

The core cytokinin signaling pathway arose early during land plant evolution and is broadly conserved (Lomin et al., 2018; Powell and Heyl, 2023). Cytokinin signaling also regulates multiple developmental processes in *M. polymorpha*, including thallus growth, meristem activity, and organ formation (Aki et al., 2019).

Functional characterization of MpMIZ1a provides further evidence for an evolutionarily conserved relationship between MIZ1 proteins and cytokinin regulation. Loss of *MpMIZ1a* affected thallus growth and hyponastic growth responses, and these phenotypes could be rescued by exogenous cytokinin treatment. In addition, genes involved in cytokinin biosynthesis were downregulated in the *Mpmiz1a* mutant, supporting a role for MpMIZ1a in maintaining cytokinin homeostasis. These findings parallel the involvement of Arabidopsis MIZ1 in cytokinin regulation (Fu et al., 2025), despite the inability of MpMIZ1a to functionally substitute for AtMIZ1. This implies that MIZ1 association with cytokinin signaling via interaction between MIZ1 and cytokinin receptors is an original, ancestral part of cytokinin signaling regulation.

What appears to have changed during evolution is not necessarily this fundamental association, but rather the molecular compatibility and developmental context in which MIZ1 proteins operate. This distinction may explain the apparent discrepancy between functional conservation within Marchantia and the lack of complementation in Arabidopsis. MpMIZ1a retains a cytokinin-related function in its native developmental context, yet cannot reproduce the specific cytokinin responses required for AtMIZ1 function when introduced into Arabidopsis. Cross-species complementation therefore reveals divergence in the implementation of MIZ1 function rather than necessarily indicating loss of its broader connection to cytokinin regulation.

### Hydrotropism may represent a derived MIZ1 function

Unlike Arabidopsis MIZ1, which is primarily studied in roots and hydrotropism, MpMIZ1a functions in developmental processes in a rootless land plant. Its strong expression in spores and rapid downregulation during germination further suggest a possible role in developmental transitions associated with reproduction and early growth. Together with its involvement in cytokinin-dependent development, these observations suggest that the ancestral function of MIZ1 proteins may have been broader than the regulation of directional root growth.

It is therefore possible that MIZ1 proteins originally participated in cytokinin-associated developmental or environmental responses and were subsequently recruited into lineage-specific developmental processes. Under this scenario, the function of AtMIZ1 in root hydrotropism would represent a derived specialization of an evolutionarily older regulatory module rather than the ancestral function of the MIZ1 family. Determining whether this model applies more broadly will require functional characterization of MIZ1 homologs in additional early-diverging land plant lineages.

Taken together, our findings support a model in which MIZ1 proteins originated early during land plant evolution as ER-associated proteins linked to cytokinin regulation by interacting with cytokinin receptors. While ER localization and the association with cytokinin receptors appear to be conserved, molecular compatibility and developmental functions have diverged between Arabidopsis and Marchantia. We propose that the role of Arabidopsis MIZ1 in hydrotropism emerged through recruitment of an evolutionarily older MIZ–cytokinin regulatory module into root-specific developmental responses.

## Material and methods

### Plant material and growth conditions

All *Arabidopsis thaliana* experiments were performed using the Columbia-0 (Col-0) ecotype as wild-type. The *miz1* loss-of-function mutant used in this study was originally described by Kobayashi et al. (2007) and was obtained from that source. Arabidopsis seedlings were grown vertically on 0.5× Murashige and Skoog (MS) medium (pH 5.9) supplemented with 1% sucrose and 0.8% agar. Plants were cultivated under long-day conditions (16 h light/8 h dark) at 22 °C, with a light intensity of approximately 250 µmol m⁻² s⁻¹.

All *Marchantia polymorpha* experiments were performed using the Takaragaike-1 (Tak-1) male and Tak-2 female accessions. Plants were cultured on half-strength Gamborg’s B5 medium containing 1% plant agar. Plants were grown in a controlled-environment chamber at 21 °C and 55% relative humidity under a 16 h light/8 h dark photoperiod, with illumination provided by white LED light at approximately 60 µmol m⁻² s⁻¹. Gametangiophore formation was induced by supplementing these light conditions with 40 µmol m⁻² s⁻¹ far-red light and the growth medium with 1% sucrose.

### Structural superposition of MIZ1 homologs

Predicted protein structures of *Arabidopsis thaliana* MIZ1 (AtMIZ1), *Marchantia polymorpha* MIZ1 (MpMIZ1), *Physcomitrium patens* MIZ1 (PpMIZ1), and *Amborella trichopoda* MIZ1 (AmTrMIZ1) were obtained from AlphaFold predictions. Structural superposition was performed in PyMOL (version 3.1, Schrödinger LLC) using the CEalign plugin. Prior to alignment, flexible low-confidence terminal regions were excluded from visualization to facilitate comparison of the conserved structural core. Pairwise structural similarity was quantified by calculating the root-mean-square deviation (RMSD) between each homolog and AtMIZ1. Structural figures were generated using PyMOL.

### Vector construction and plant transformation

Coding sequences (CDSs) of *MIZ1*-like genes were amplified from representative land plant species using gene-specific primers. The following genes were used: *MpMIZ1a* (*Marchantia polymorpha*; Mp2g13620.1), *MpMIZ1b* (*M. polymorpha*; Mp3g00040), *PpMIZ1c* (*Physcomitrium patens*; Pp3c19_15840V3.3), *PpMIZ1d* (*P. patens*; Pp3c18_14150V3.1), *SmMIZ1* (*Selaginella moellendorffii*; Sm91391), and *AmTrMIZ1* (*Amborella trichopoda*; AmTr_v1.0_scaffold00053.180).

All CDSs were first cloned into the Gateway entry vector *pDONR221* using BP recombination reactions. For subcellular localization and cross-species complementation analyses in *Arabidopsis thaliana*, the CDSs were transferred into the Gateway destination vector *pB7FWG2*, generating C-terminal GFP fusion constructs under the control of the CaMV 35S promoter. Transgenic Arabidopsis plants were selected on Basta-containing medium, and GFP fluorescence was verified by confocal laser scanning microscopy.

For the Arabidopsis MIZ1 control, the *AtMIZ1-GFP* construct was generated following the approach described by Yamazaki et al. (2012) and cloned into the Gateway destination vector *pK7WG2*. Transgenic lines were selected on kanamycin-containing medium, and GFP fluorescence was confirmed by confocal microscopy. All Arabidopsis transformations were performed using *Agrobacterium tumefaciens*-mediated floral dip. Independent transgenic lines were selected and propagated for further analyses.

To generate Marchantia CRISPR mutants, guide RNAs were designed using the CasFinder tool. Annealed guide RNA oligonucleotides were cloned into the *pMpGE_En03* entry vector at the *BsaI* site using T4 DNA ligase (NEB; Cat. No. M0202S), as described by Sugano et al. (2018). Gateway LR Clonase enzyme (Thermo Fisher Scientific; Cat. No. 1791020) was used to transfer the guide RNA and CRISPR cassette from the *pMpGE_En03* entry vector into the Gateway destination vector *pMpGWB101*. For translational reporter lines, constructs were generated using either a 2.1-kb *MpMIZ1* promoter region including the 5′ UTR or the ubiquitin 2 promoter (*pUBE2*) to drive expression of the *MpMIZ1* CDS fused to an mNeonGreen (mNG) tag. The resulting fragments were cloned into *pMpGWB301* digested with *HindIII* and *SacI* using the Gibson Assembly kit (NEB; Cat. No. E5510S).

### Hydrotropism assay

Root hydrotropic responses were analyzed using a split-agar system as previously described (Eapen et al., 2015), with minor modifications. Briefly, water stress medium (WSM) consisted of half-strength Murashige and Skoog (MS) basal salts supplemented with 0.5% (w/v) sucrose, 0.9% (w/v) agar, 2.5% (v/v) glycerol, and 0.5% (w/v) alginic acid. The pH of the medium was adjusted to 5.7 prior to the addition of alginic acid and agar. All media preparation and plate assembly were performed under sterile conditions in a laminar flow hood.

The assay system consisted of Petri dishes containing two horizontally arranged medium sectors: normal medium (NM), lacking osmotic additives, and WSM. Seedlings were positioned at the interface between the two media to allow directional root growth in response to the imposed water potential gradient.

Plates were incubated vertically under standard growth conditions, and hydrotropic responses were documented by digital imaging at the indicated time points. Primary root length and bending angles were quantified from digital images using ImageJ.

### Cytokinin (zeatin) treatment

For cytokinin sensitivity assays, seedlings were grown on half-strength Murashige and Skoog (MS) medium supplemented with 2 µM trans-zeatin. Trans-zeatin was prepared as a stock solution in DMSO and added to autoclaved medium after cooling to approximately 55 °C. Control medium contained an equivalent volume of DMSO without trans-zeatin.

Seeds were sown directly onto trans-zeatin-containing or control medium and grown vertically under standard growth conditions. Primary root growth was monitored at the indicated time points, and root length was quantified from digital images using ImageJ.

### Phylogenetic analysis of MIZ1 homologs

MIZ1 protein homologs were identified using HMMER v3.3 searches against the NCBI non-redundant (nr) protein database, using the *Arabidopsis thaliana* MIZ1 protein (AT2G41660) as the query. To reduce sequence redundancy, retrieved sequences were clustered at 45% amino acid identity using USEARCH.

The resulting non-redundant protein sequences were aligned using the ClustalW algorithm implemented in MEGA. Phylogenetic relationships were inferred using three independent methods: neighbor-joining (NJ), maximum parsimony (MP), and maximum likelihood (ML). NJ trees were constructed using Poisson-corrected distances with pairwise deletion of gaps. MP analysis was performed using default parameters. For ML analysis, the best-fitting amino acid substitution model was determined prior to tree reconstruction.

The robustness of tree topologies was evaluated by bootstrap analysis with 1,000 replicates for all phylogenetic methods. All MIZ1-like sequences included in the analysis are listed in Supplementary Table 1, and the resulting phylogenetic tree is shown in Figure 1C.

### Bimolecular fluorescence complementation in *N. benthamiana* leaves

Bimolecular fluorescence complementation (BiFC) assays were performed by transient expression in *Nicotiana benthamiana* leaves. Coding sequences of the proteins of interest were fused in-frame to the N- or C-terminal fragment of YFP and introduced into *Agrobacterium tumefaciens* strain GV3101. Agrobacterial cultures carrying the corresponding construct pairs were mixed at equal ratios and infiltrated into the abaxial side of leaves from 4–6-week-old plants. Reconstituted fluorescence was analyzed 2–3 d after infiltration using an LSM 910 laser-scanning confocal microscope (Carl Zeiss Microscopy). Images were acquired using identical laser power, detector gain, and emission settings for all samples within an experiment. The specificity of the BiFC signal was assessed using the corresponding negative-control combinations. Each experiment was repeated independently at least three times using leaves from different plants.

### Arabidopsis suspension culture maintenance

*Arabidopsis thaliana* Col-0 root-derived suspension cells were maintained using modified procedures based on Blackhall (1992) and Mathur and Koncz (1998). Cells were cultured in liquid medium containing 4.32 g L⁻¹ Murashige and Skoog (MS) basal medium without vitamins (Duchefa, M0221), 30 g L⁻¹ sucrose (Sigma-Aldrich), 0.25 mg L⁻¹ 2,4-dichlorophenoxyacetic acid (2,4-D; Duchefa, D091), 0.015 mg L⁻¹ kinetin (Sigma-Aldrich, K0753), and 2 mL L⁻¹ B5 vitamin stock (1 g L⁻¹ nicotinic acid, 1 g L⁻¹ pyridoxine HCl, 10 g L⁻¹ thiamine HCl, and 100 g L⁻¹ myo-inositol). The medium was adjusted to pH 5.7 with KOH, filtered, and subsequently sterilized by autoclaving. Cultures were maintained in 250-mL Erlenmeyer flasks containing 50 mL of culture at 20–22 °C with continuous shaking at approximately 120 rpm and were subcultured weekly at a 1:10 dilution.

### Protoplast isolation

Protoplast isolation was performed using a modified procedure based on Mathur and Koncz (1998). All centrifugation steps were performed at room temperature using reduced acceleration and deceleration settings. Protoplasts were isolated from 2–3-day-old suspension cultures. A 50-mL suspension culture was centrifuged for 5 min at 480 × *g*, and the cells were resuspended in enzyme solution containing 1% (w/v) cellulase Onozuka RS (Duchefa) and 0.2% (w/v) macerozyme (Yakult) in glucose–mannitol (GM) buffer (4.4 g L⁻¹ MS basal salts, 0.17 M glucose, and 0.17 M mannitol; pH 5.5, adjusted with KOH). The suspension was incubated for 3–4 h in the dark at room temperature with gentle shaking.

Following digestion, protoplasts were centrifuged for 5 min at 310 × *g*, gently resuspended in approximately 5 mL of GM buffer, and diluted to a final volume of 25 mL with GM buffer. Protoplasts were washed by centrifugation for 5 min at 215 × *g* and resuspended in 5 mL of S buffer (4.4 g L⁻¹ MS basal salts and 0.28 M sucrose; pH 5.5, adjusted with KOH). The suspension was centrifuged for 7 min at 140 × *g*, and protoplasts floating at the surface were collected and adjusted to a final concentration of 4 × 10⁶ cells mL⁻¹ with GM buffer.

### PEG-mediated protoplast transfection

Protoplast transfection was performed using a modified procedure based on Mathur and Koncz (1998). Plasmid DNA was purified using the E.Z.N.A. Plasmid DNA Maxi Kit (Omega Bio-tek, D6922-00S) and concentrated by isopropanol precipitation to approximately 1 µg µL⁻¹. For each transfection, 3 µg of each plasmid was combined in a total volume of 15 µL and mixed with 50 µL of protoplast suspension. Subsequently, 150 µL of PEG solution containing 25% (w/v) PEG 6000, 0.45 M mannitol, and 0.1 M Ca(NO₃)₂ (pH 9.0, adjusted with KOH) was added. Samples were incubated for 1 h in the dark, after which the PEG solution was diluted by two sequential additions of 0.5 mL of 0.275 M Ca(NO₃)₂, with gentle mixing after each addition. Protoplasts were centrifuged for 7 min at 150 × *g*, resuspended in 0.5 mL of GM buffer, and incubated overnight in the dark before imaging.

### Confocal imaging and colocalization analysis

Confocal imaging was performed using a Leica SP8 confocal laser-scanning microscope equipped with HC PL APO 20×/0.75 CS dry, HC PL APO 40×/1.10 W CORR CS2 water-immersion, and HC PL APO 63×/1.30 CORR CS glycerol-immersion objectives. GFP and mCherry were excited using 488- and 561-nm laser lines, respectively. Acquisition settings were selected to minimize spectral bleed-through between channels and were kept constant among samples compared within the same experiment.

Colocalization was analyzed on maximum-intensity projections in Fiji using the Coloc 2 plugin. Colocalization was quantified using the thresholded Manders’ coefficient tM1 following automatic threshold determination using the Costes method. tM1 represents the fraction of the GFP signal overlapping with the mCherry-HDEL signal. Analyses were performed on the full images without applying an additional region-of-interest mask.

### Confocal microscopy

Confocal imaging of *Arabidopsis thaliana* seedlings was performed using a Zeiss LSM 910 inverted confocal laser-scanning microscope. GFP was excited using the 488-nm laser line, and fluorescence emission was detected between 500 and 550 nm. For experiments involving the endoplasmic reticulum marker, mCherry was excited using the 561-nm laser line, and fluorescence emission was detected between 580 and 630 nm. All images were acquired using identical laser power, detector gain, and pinhole settings for samples compared within each experiment. Image processing and analysis were performed using Zeiss ZEN software or ImageJ, as indicated.

Marchantia gemmae and spores were imaged using a Leica STELLARIS 5 confocal microscope equipped with hybrid detectors and a pulsed white-light laser. mNeonGreen was excited at 506 nm, and fluorescence emission was detected between 512 and 560 nm. Laser output was set to 8%, and detectors were operated in photon-counting mode. Time gating was applied to suppress chlorophyll autofluorescence. Z-stacks were acquired using a 20× dry objective, and maximum-intensity projections were generated using ImageJ.

### Protein structure prediction and protein-protein docking

Protein structure predictions were generated using AlphaFold2 implemented through ColabFold (Jumper et al., 2021; Mirdita et al., 2022). Amino acid sequences of AtMIZ1, AtAHK2, MpMIZ1a, and MpCHK were submitted individually, and the resulting predicted structures were used for subsequent analyses. Structural models with the highest confidence scores were selected for protein–protein docking.

Protein–protein docking was performed using HDOCK, which combines template-based modeling and *ab initio* docking approaches. Docking analyses were performed between AtMIZ1 and AtAHK2 and between MpMIZ1a and MpCHK. The top-ranked docking models generated by HDOCK were selected for further analysis.

Predicted protein complexes were visualized using PyMOL (Schrödinger, LLC). Putative interaction interfaces were identified by examining residues located at intermolecular distances of 2.5–3.0 Å. Potential hydrogen bonds and salt bridges were manually inspected and annotated based on the predicted docking models. Interface residues identified in the predicted complexes were visualized as stick representations and used for structural comparison between the Arabidopsis and Marchantia protein complexes.

### RNA extraction and RT-qPCR analysis

Total RNA was extracted from *Marchantia polymorpha* Tak-1, *miz1a#1*, *miz1a#2*, *miz1b#12*, *miz1b#13*, *pUBE2::MIZ1A-mNG*, and *pUBE2::MIZ1B-mNG* plants grown under mock conditions or supplemented with 50 μM *N*6-isopentenyladenine (iP). RNA was extracted using the RNeasy Plant Mini Kit (Qiagen), followed by DNase I treatment (Thermo Fisher Scientific) to remove genomic DNA contamination. cDNA synthesis was performed using the iScript cDNA Synthesis Kit (Bio-Rad) according to the manufacturer’s instructions.

Quantitative reverse transcription PCR (RT-qPCR) was performed using SYBR Green Supermix (Bio-Rad) on a Roche LightCycler 480 Real-Time PCR System. Reaction setup was automated using a Beckman Coulter Biomek i5 liquid-handling robot. Previously published gene-specific primers were used for *MpLOG1*, *MpIPT1*, *MpRRA*, and *MpRRB*, with *MpEF1α* used as the reference gene. Relative transcript levels were calculated using the ΔΔCt method and expressed relative to Tak-1 within the corresponding treatment. Melt-curve analysis was performed to verify amplification specificity.

## Author contributions

A.S., S.D., and J.F. conceived the study. A.S. performed the Arabidopsis experiments, analyzed the data, and wrote the manuscript. S.D. performed the Marchantia experiments and contributed to writing the manuscript. D.V. and M.T. performed the BiFC experiments. A.T. performed the protoplast co-localization experiments. M.L. performed the RT-qPCR analyses. E.B. and J.F. supervised the study and revised the manuscript. All authors approved the final manuscript.

## Acknowledgements

We thank the Imaging and Optics Facility at the Institute of Science and Technology Austria (ISTA) for access to microscopy infrastructure and technical support. We are grateful to members of the Friml laboratory for discussions and feedback throughout this work.

