## Supporting Information for "Conserved molecular compatibility for MIZ1 function in cytokinin signaling in land plants"

**Supplementary information:**

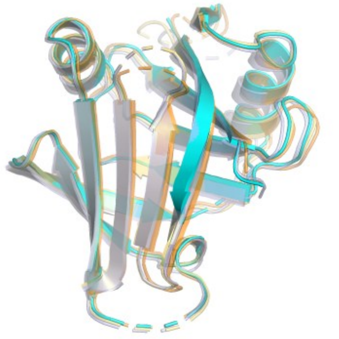

**Figure S1. Structural conservation of MIZ1 homologs across land plants. (A)** Structural superposition of predicted MIZ1 proteins from *Arabidopsis thaliana* (AtMIZ1, **grey**), *Amborella trichopoda* (AmTrMIZ1, **light blue**), *Physcomitrium patens* (PpMIZ1, **orange**), and *Marchantia polymorpha* (MpMIZ1, **yellow**). Predicted structures were aligned using CEalign in PyMOL, revealing a highly conserved overall protein fold despite evolutionary divergence. **(B)** Root-mean-square deviation (RMSD) values for pairwise structural alignments of MIZ1 homologs relative to AtMIZ1.

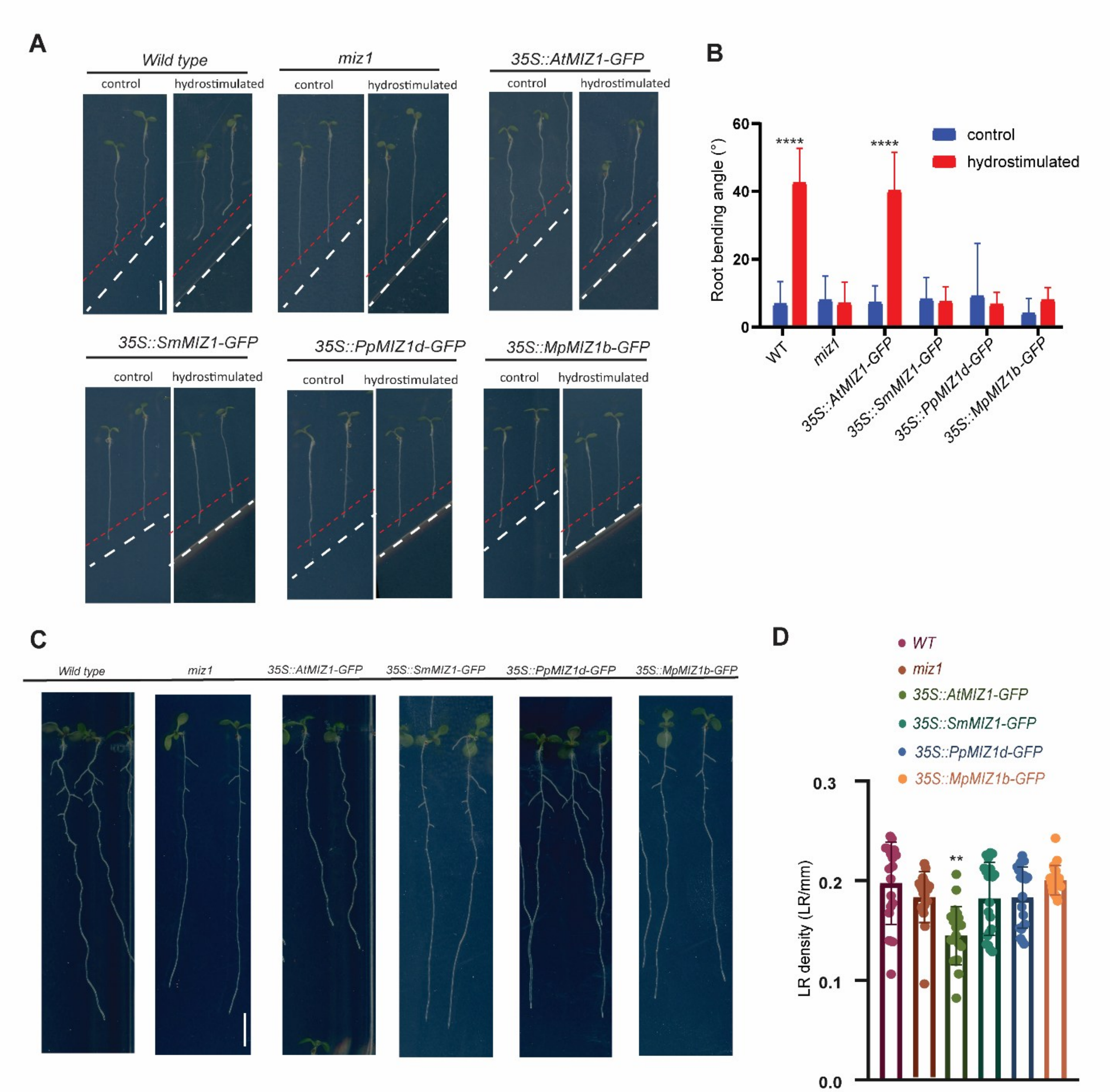


**Figure S2. Hydrotropic response and lateral root development in transgenic *Arabidopsis* lines expressing MIZ1 homologs.**

(A) Representative images of the hydrotropic response of wild-type (WT), *miz1*, *35S::AtMIZ1-GFP, 35S::SmMIZ1-GFP, 35S::PpMIZ1D-GFP*, and *35S::MpMIZ1B-GFP* seedlings grown under hydrostimulation conditions. (B) Quantification of the hydrotropic response shown in (A). Data are presented as bending angle (°) or hydrotropic response index, as indicated. (C) Representative images of lateral root development in WT, *MIZ1*, *35S::AtMIZ1-GFP, 35S::SmMIZ1-GFP, 35S::PpMIZ1D-GFP*, and *35S::MpMIZ1B-GFP* seedlings. (D) Quantification of lateral root density shown in (C). Data are expressed as lateral roots per millimeter of primary root length (LR/mm). Statistical analyses were performed as indicated.

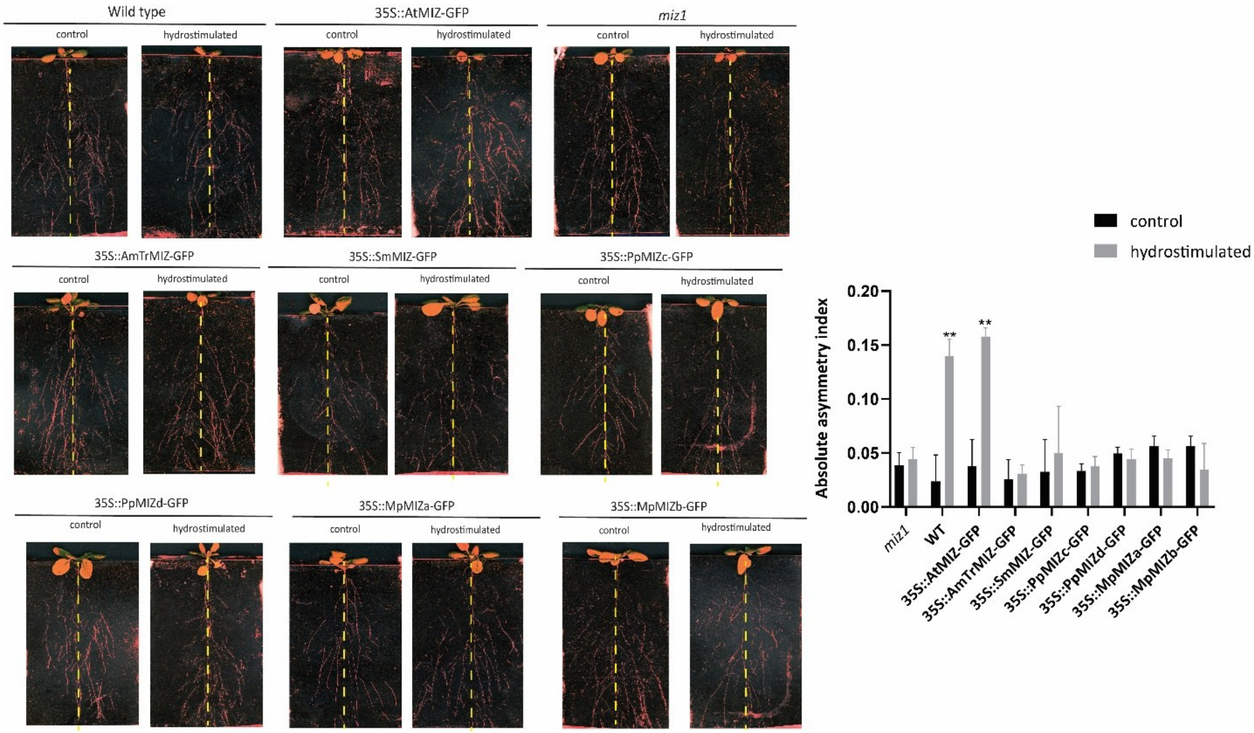


**Figure S3. Hydrotropic responses of transgenic lines in rhizotrons.** (A) Representative images of transgenic lines grown in soil-filled rhizotrons containing a hydrostimulated chamber and a control chamber. Roots were allowed to grow between the two compartments and their growth direction was monitored over time. (B) Absolute asymmetry index of the transgenic lines grown in the rhizotron assay. The asymmetry index was calculated based on root distribution between the hydrostimulated and control chambers. Data are shown as mean ± SEM. Statistical significance was determined as indicated.

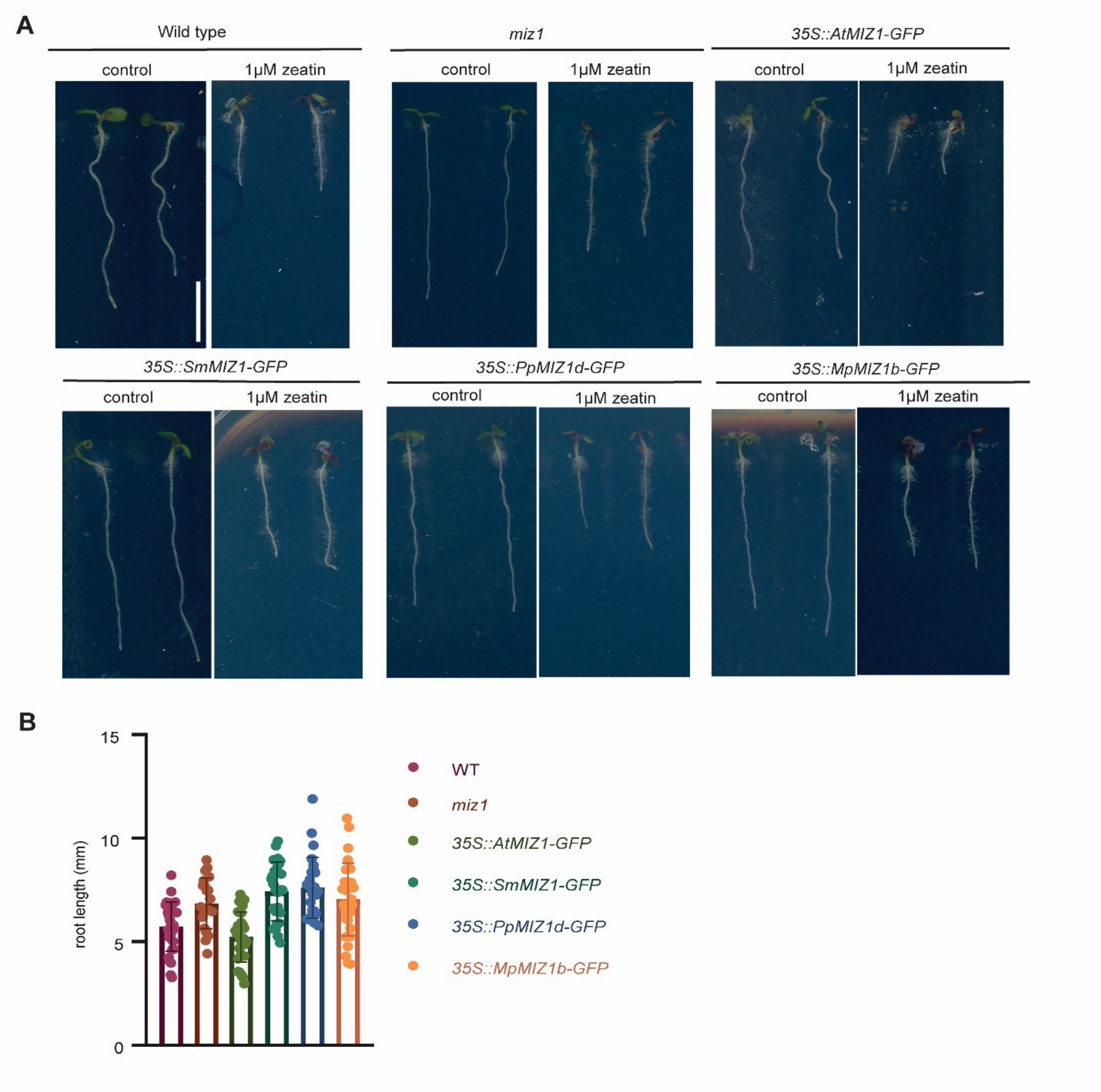


**Figure S4. Response of transgenic *Arabidopsis* lines expressing MIZ1 homologs to zeatin treatment.**
(A) Representative images of WT*, miz1, 35S::AtMIZ1-GFP, 35S::SmMIZ1-GFP, 35S::PpMIZ1D-GFP*, and *35S::MpMIZ1B-GFP* seedlings grown in the presence of zeatin.
(B) Quantification of the zeatin response shown in (A). Data are presented as mean ± SEM. Statistical analyses were performed as indicated.

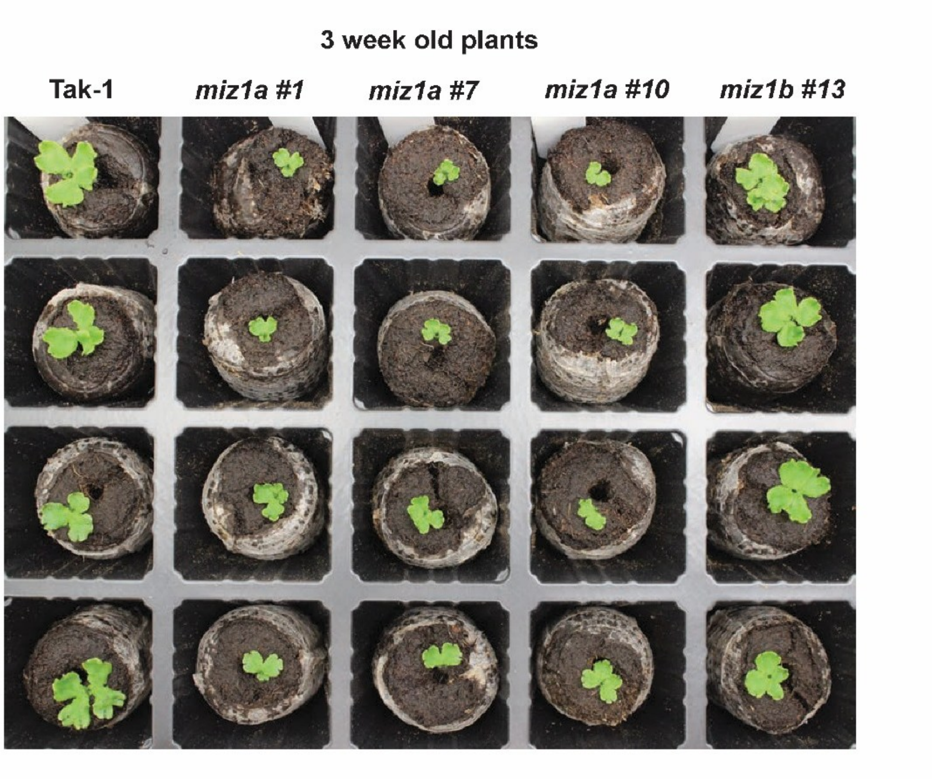


**Figure S5. Growth phenotypes of *Marchantia polymorpha miz1* mutants on soil.** Representative images of 3-week-old *Marchantia polymorpha* plants grown in soil. Genotypes include *miz1a* mutant lines (#1 and #7), *miz1b* mutant lines (#10 and #13).

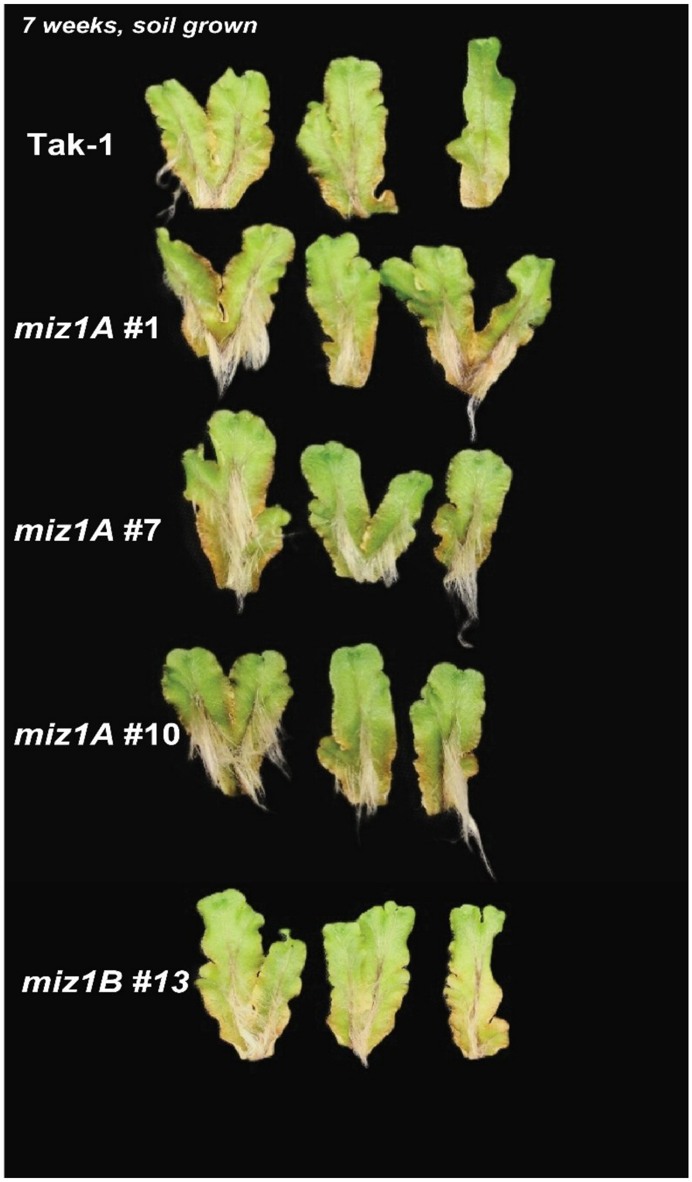


**Figure S6. Rhizoid development in transgenic *Marchantia polymorpha* lines.** Representative images of 7-week-old *Marchantia polymorpha* thalli grown in soil. Rhizoid development is shown for the indicated transgenic lines and mutants.

–

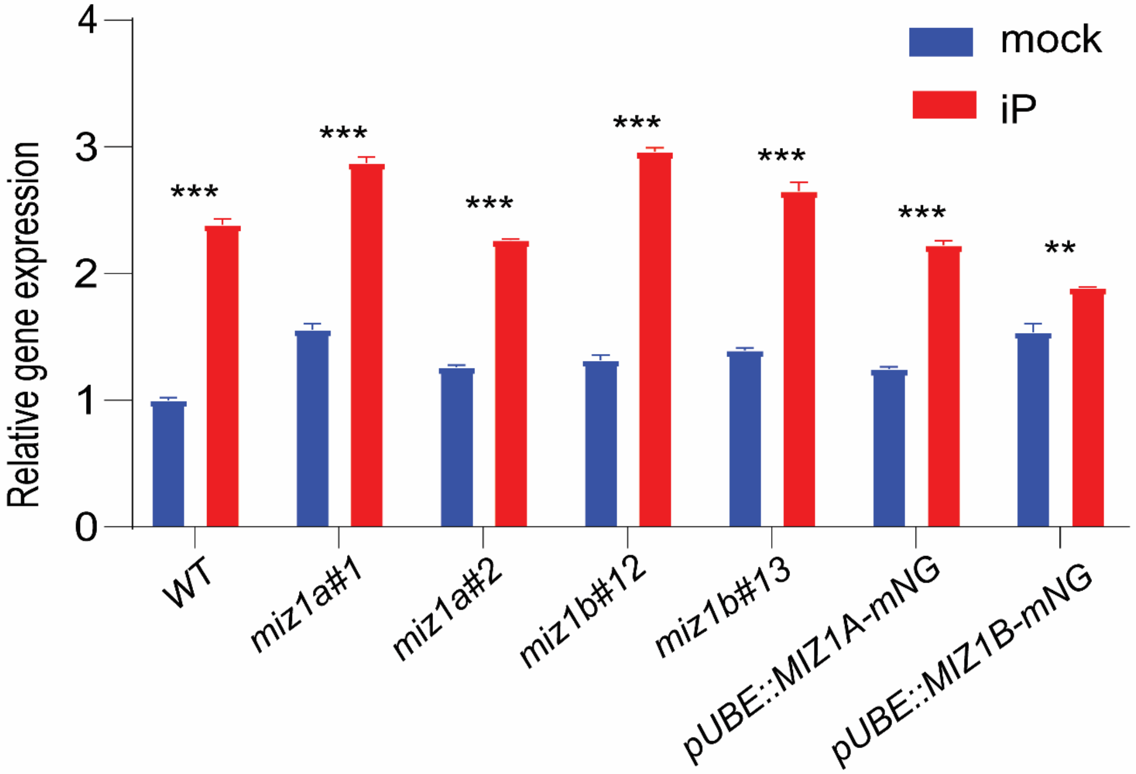


**Figure S7. Relative expression of cytokinin-responsive genes in *Marchantia polymorpha* MIZ1 lines.** Relative *MpRRA* expression in Tak1, *miz1a*#1, *miz1a*#2, *miz1b*#12, *miz1b*#13, pUBE2::*MIZ1a*-mNG, and pUBE2::*MIZ1a*-mNG plants grown under mock or 50 µM iP conditions, as determined by quantitative RT-qPCR. Transcript levels were normalized to the reference gene and expressed relative to Tak-1 under mock conditions. Statistical comparisons between each genotype and Tak-1 within the corresponding treatment were performed using two-way ANOVA followed by Sidak's multiple-comparisons test. Bars represent the mean ± SD, and individual points represent technical replicates. **P** < 0.01 (**); **P** < 0.0001 (****).

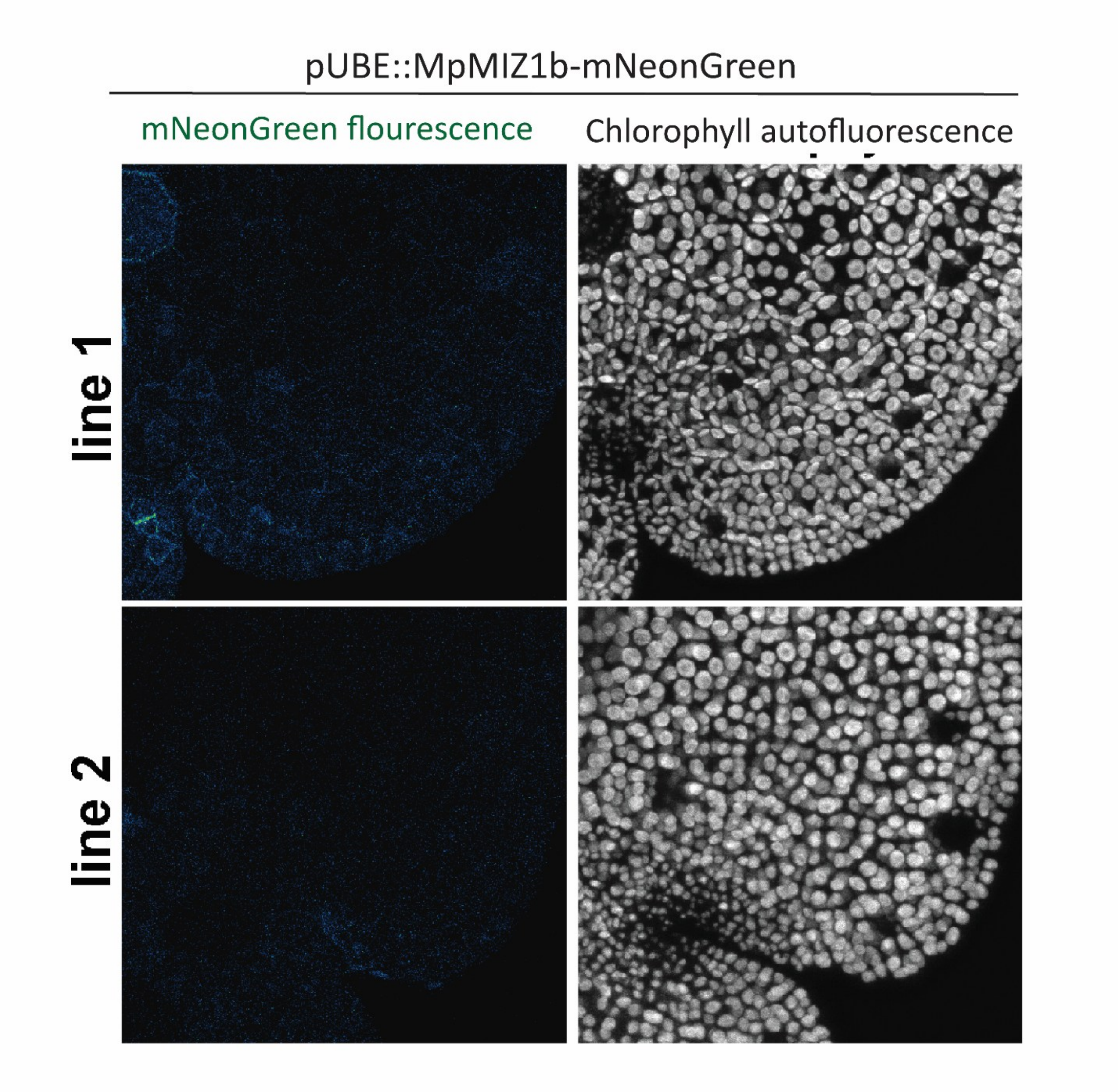


**Figure S8. Expression of *pUBE2::MIZ1b-mNeonGreen* in *Marchantia polymorpha* gemmae.** Confocal microscopy images of *pUBE2::MIZ1b-mNeonGreen* plants showing the absence of detectable mNeonGreen signal in gemmae

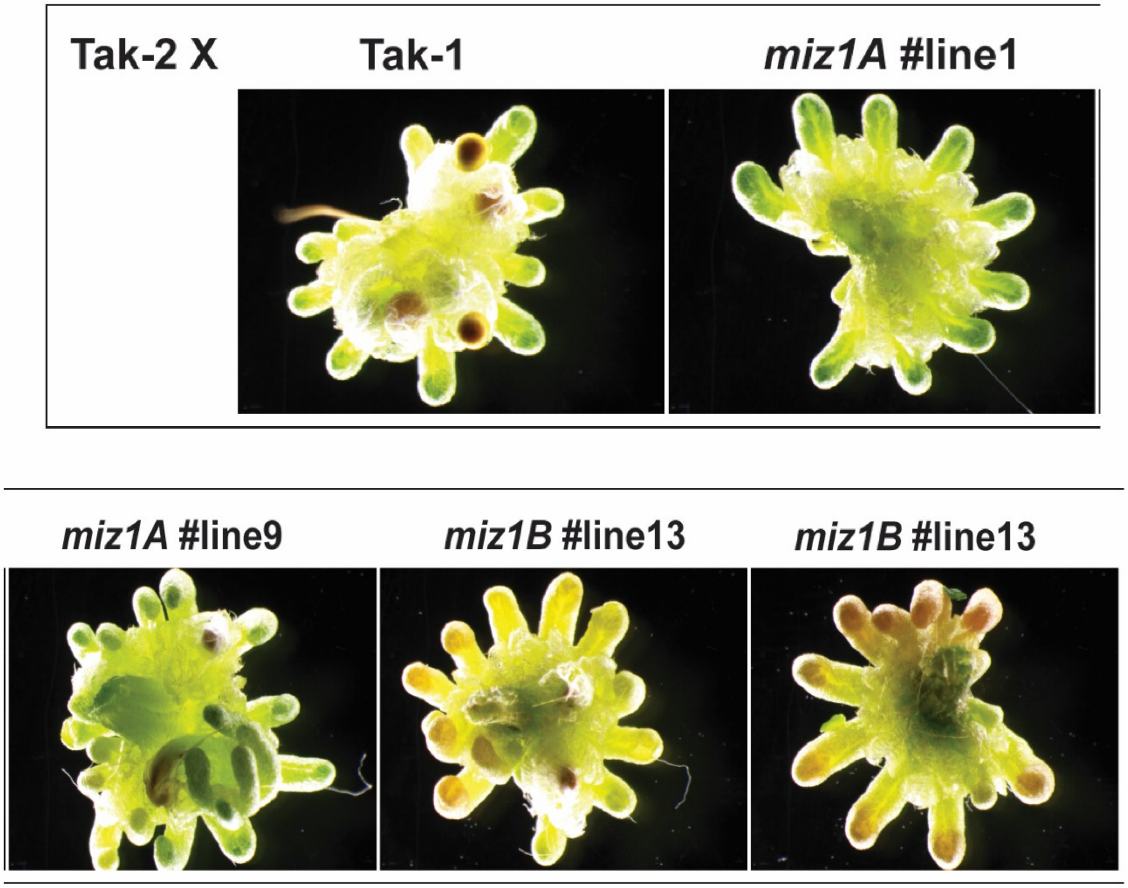


**Figure S9. Sporophyte development in *Marchantia polymorpha* transgenic lines.** Representative images of sporophytes produced by *Marchantia polymorpha* transgenic lines, including *miz1a* mutant lines (#1 and #9), *miz1b* mutant lines (#13 and #14), and *pUBE2::MIZ1a* and *pUBE2::MIZ1b* overexpression lines.

| Comparison | RMSD |
|---|---|
| **AtMIZ1 – MpMIZ1a** | **1.83 Å** |
| **AtMIZ1 – AmTrMIZ1** | **2.66 Å** |
| **AtMIZ1 – PpMIZ1** | **2.86 Å** |

**Supplementary Table 1. Structural conservation of MIZ1 homologs across land plants.**

**(A)** Structural superposition of predicted MIZ1 proteins from *Arabidopsis thaliana* (AtMIZ1), *Amborella trichopoda* (AmTrMIZ1), *Physcomitrium patens* (PpMIZ1), and *Marchantia polymorpha* (MpMIZ1). Predicted structures were aligned using CEalign in PyMOL, revealing a highly conserved overall protein
